# Functional reorganization of motor cortex connectivity during learning

**DOI:** 10.64898/2026.03.03.709199

**Authors:** Kayvon Daie, Kyle Aitken, Márton Rózsa, Matthew S. Bull, Peter C. Humphreys, Ziheng Christina Wang, Lucas Kinsey, Mohit Kulkarni, Kimberly L. Stachenfeld, Maria K. Eckstein, Zeb Kurth-Nelson, Claudia Clopath, Timothy P. Lillicrap, Matthew Botvinick, Matthew Golub, Stefan Mihalas, Karel Svoboda

**Author notes:** Equal contributors (Kayvon Daie, Kyle Aitken).

## Abstract

Learning new tasks requires the brain to reshape the flow of neural activity, but how these changes arise from dynamic neural connectivity remains unclear. Here, we used two-photon photostimulation and calcium imaging to map learning-related changes in connectivity in layer 2/3 of mouse motor cortex, induced by learning of an optical brain-computer interface (BCI) task. Mice rapidly (within minutes) learned to change activity in a conditioned neuron to earn rewards. Activity changes were sparse; the conditioned neuron increased activity more than surrounding neurons. Mapping connectivity before and after learning revealed changes in motor cortex connectivity, enriched in neurons that were active before trial initiation, analogous to motor cortex populations that are active preceding movement. Motor cortex plasticity reroutes preparatory activity to neurons that are active later and control the conditioned neuron. Our findings show how rapid learning can be achieved through structured changes in motor cortex connectivity.

## Introduction

Voluntary movements are generated by dynamic neural activity in motor cortex (Evarts 1968). At the level of neural populations, distinct movements are associated with their own neural trajectories (Georgopoulos et al. 1982; Li et al. 2016; Gallego et al. 2017; Elsayed et al. 2016). Movement-related trajectories are preceded by preparatory activity that emerges before movement onset and sets the initial conditions from which the subsequent movement-generating dynamics unfold (Churchland et al. 2006; Shenoy et al. 2011; Churchland et al. 2012). These dynamics are shaped by both recurrent circuits within motor cortex (Daie et al. 2021; Finkelstein et al. 2025) and long-range inputs (Guo et al. 2017; Sauerbrei et al. 2020; Perich et al. 2018), providing a multi-regional substrate that can be reshaped to support learning.

During learning, population-level motor cortex activity reorganizes in systematic ways (Huber et al. 2012; Peters et al. 2014; Sadtler et al. 2014), including changes in neural trajectories and preparatory activity (Perich et al. 2018; Vyas et al. 2018; Sun et al. 2022). These learning-related changes in neural dynamics are widely thought to arise from synaptic plasticity, but where in the brain this plasticity occurs, and how it reshapes motor cortical dynamics, remains unclear.

During rapid learning (< 1 hour), the low-dimensional structure of motor cortex activity, including correlations, is largely preserved, despite changes in neural trajectories. This has led to the suggestion that motor cortex connectivity is stable and rapid learning occurs mainly in upstream brain areas (Sadtler et al. 2014; Golub et al. 2018; Menéndez et al. 2025). However, cortical neurons can express synaptic plasticity over times of seconds *in vitro* (Markram et al. 1997; Sjöström et al. 2001; Froemke et al. 2010) and *in vivo* (Bittner et al. 2017), consistent with rapid learning.

Moreover, for slower motor learning (days), synaptic changes have been detected in the motor cortex (Karni et al. 1998; Rioult-Pedotti et al. 2000; Holtmaat et al. 2005; Peters et al. 2017), in addition to brain regions providing input to the motor cortex (Yin et al. 2009; Audette et al. 2019). However, these studies could not resolve whether plasticity occurred between the neurons whose activity reorganized during learning. How motor cortex plasticity reshapes movement-related dynamics during rapid learning remains unresolved.

A major barrier to linking synaptic plasticity and learning is the difficulty of measuring changes in connectivity in the learning brain. Two-photon optogenetics enables targeted photostimulation of defined neurons while simultaneously recording activity across the local circuit using calcium imaging (Rickgauer et al. 2014; Packer et al. 2015). By quantifying photostimulus-evoked changes in neuronal activity, this approach enables mapping of ‘causal connectivity’—the directed influence of photostimulating a defined neuronal population on the activity of other neurons in the local network (Chettih & Harvey 2019; Daie et al. 2021; Oldenburg et al. 2024; Finkelstein et al. 2025).

In naturalistic motor learning, behavior-related changes in neural activity are distributed across multiple brain regions, making it difficult to localize where learning-related plasticity is expressed and which local connections should be examined. In brain–computer interfaces (BCIs), learning must involve changes in the activity of experimenter-defined groups of ‘conditioned neurons’ (CNs) (Fetz 1969). Conditioned neurons are by construction a key locus of learning. By combining mapping of causal connectivity with BCI learning, we examined learning-related changes in activity and changes in connectivity within the same population of motor cortex neurons.

We developed an optical BCI paradigm for motor cortex (MC) neurons (Clancy et al. 2014; Prsa et al. 2017). Calcium imaging allowed us to track activity in the same neuronal population over weeks as mice learned to control a BCI using different individual neurons across daily sessions. Between sessions, we measured the causal connectivity using two-photon photostimulation. Neurons whose activity changed during learning also exhibited systematic changes in local connectivity, with the strongest modifications occurring in neurons participating in preparatory activity. Network modeling and population analyses revealed that these connectivity changes are explained by local recurrent plasticity, which reconfigures preparatory activity to drive BCI-relevant neural patterns. Together, these results provide direct evidence that rapid learning is supported by plasticity in local motor cortical circuits that selectively modifies neurons encoding preparatory signals.

## Results

### An optical brain-computer interface task

Mice were trained to control a brain-computer interface (BCI) using the activity of a single conditioned neuron (CN) in layer 2/3 of the primary motor cortex (MC; Fig. 1a). Population activity was simultaneously recorded using two-photon calcium imaging across a 1 mm × 1 mm field-of-view (FoV; median: 481 neurons; 1st–3rd quartile: 364–622). When CN activity exceeded a threshold, it was converted into a control signal that moved a motorized reward port towards the mouse (Fig. 1a).

**Figure 1:**
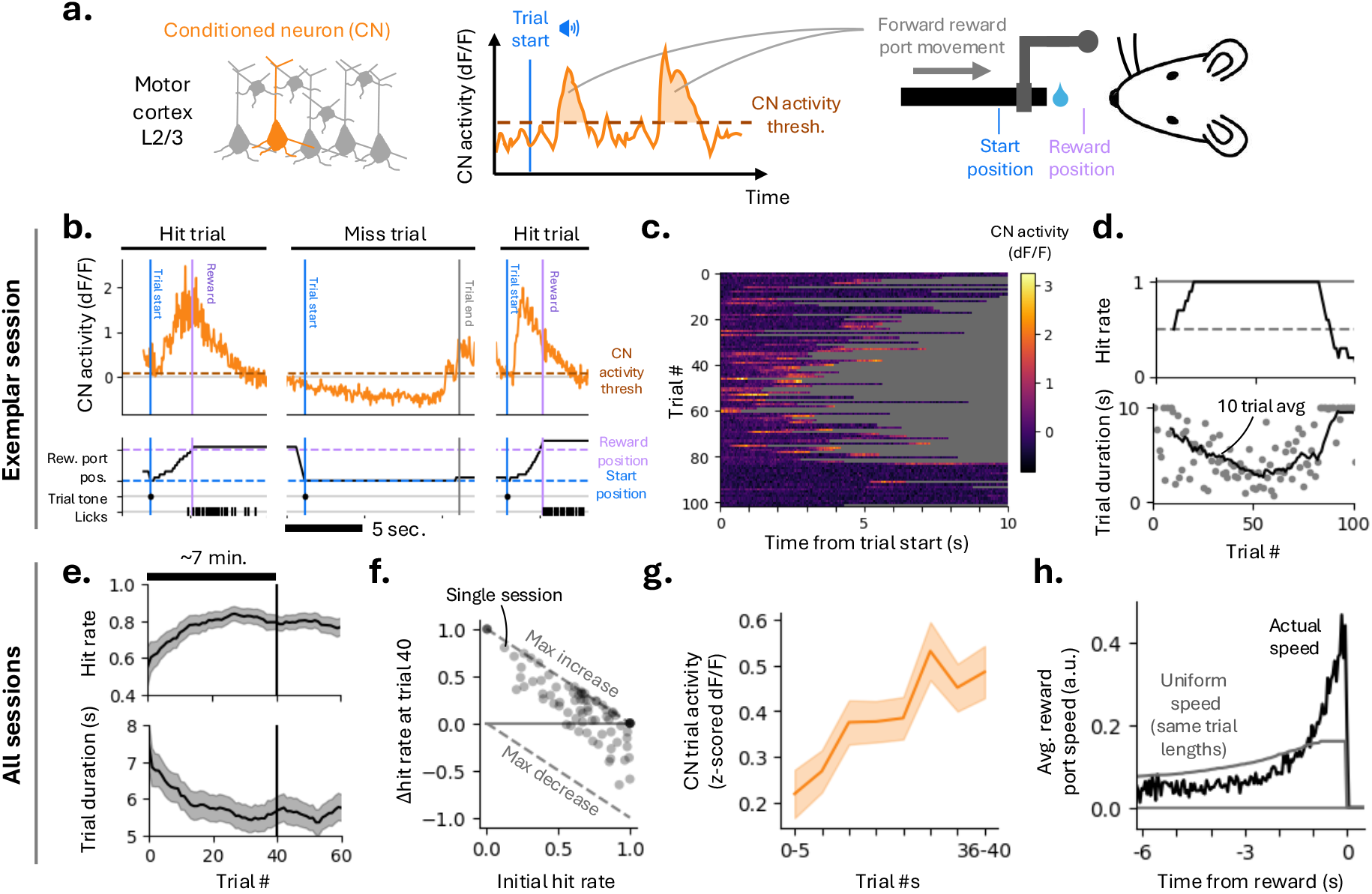
Rapid learning of a single-neuron brain-computer interface. **(a)** BCI task. Activity of a single Layer 2/3 pyramidal neuron (the ‘conditioned neuron’; CN) in the primary motor cortex controls reward port speed. **[b-d]** Exemplar session. **(b)** Three consecutive trials (hit, miss, hit). CN activity (top) drives reward port movement (bottom). **(c)** CN activity across trials within the session. **(d)** Hit rate (top; running average over 10 trials) and trial duration (bottom). The decrease in hit rate at the end of the session reflects animal disengagement. **[e-h]** Aggregate data (78 sessions, 10 mice). **(e)** Mean hit rate (top) and trial duration (bottom) as a function of trial # within a session (mean across sessions; shaded area, 95% CI from bootstrap). The vertical black line marks trial 40, which was used as a uniform cutoff across sessions and mice to restrict analyses to the early learning period during which robust performance improvements were observed (median duration ≈7 min). **(f)** Change in hit rate (Δ hit rate) versus initial hit rate for each session. **(g)** CN trial activity as a function of trial #, shown in 5-trial bins. CN activity was *z*-scored within each session before pooling (mean ± s.e.m.). **(h)** Mean reward port speed as a function of time to reward, showing a pronounced increase late in the trial preceding reward delivery. The grey line indicates the expected speed if port movement were uniform throughout each trial, with trial durations fixed.

Each trial began with an auditory cue signaling trial onset, with the reward port initially positioned beyond the reach of the mouse. A pole attached to the reward port provided real-time information about proximity to reward through whisker-based somatosensation. Trials were rewarded if CN activity brought the port within reach of the tongue (reward position) within 10 s (hit trial, Fig. 1b); otherwise, the trial was terminated and the port returned to the start position (miss trial). New trials were initiated only after CN activity fell below threshold following reward delivery.

Each field-of-view (24 FoV across 10 mice) was imaged for multiple days (median: 3 days; range: 1-8 days), enabling longitudinal measurement of task-related activity in the same neuronal population across sessions. At the start of each session, a new CN within the same FoV was selected. Many neurons exhibited transient modulations following the auditory trial-start cue that could already support favorable BCI task performance. CN neurons were therefore selected to have (1) weak trial-start tuning, requiring changes in CN activity to acquire reward consistently, and (2) substantial activity, as silent neurons never lead to reward port movement and thus do not provide feedback for learning to the mouse (Methods).

### BCI learning is rapid

CN activity increased across trials within a session (Fig. 1c), accompanied by a higher proportion of rewarded trials (hits) and shorter trial durations (Fig. 1d). Consistent with this increase in CN activity, hit rates rose significantly over tens of trials (Figs. 1e-f; mean increase at trial 40: 23%, 16–31%, 95% CI). Reward rate also increased over the first 40 trials of training (mean: 0.62 rewards per minute, 0.22–1.03, 95% CI), together with a reduction in trial duration (Fig. 1e; mean reduction: 0.90 s, 0.39–1.42, 95% CI).

Session-averaged CN response amplitudes increased alongside task performance (Fig. 1g). By trial 40, the mean CN *z*-scored Δ*F*/*F* amplitude had increased 2.2-fold relative to the session start (from 0.22 to 0.49; mean change, 0.27; 0.09–0.44, 95% CI). Reward port movements were driven by brief bursts of CN activity, resulting in rapid reward port displacement rather than gradual, stepwise movement throughout the trial (Fig. 1b-c). The majority of movement occurred within one second preceding reward (Fig. 1h).

### Changes in neural activity are sparse

During BCI learning, the mouse must infer the identity of the CN through feedback arising from its influence on reward port movement. One strategy is to infer and selectively modulate the CN (‘sparse’; Fig. 2a). Alternatively, activity could be modulated across a large fraction of neurons (‘dense’). These scenarios represent two extremes along a continuum defined by how selectively CN activity changes relative to the surrounding population.

**Figure 2:**
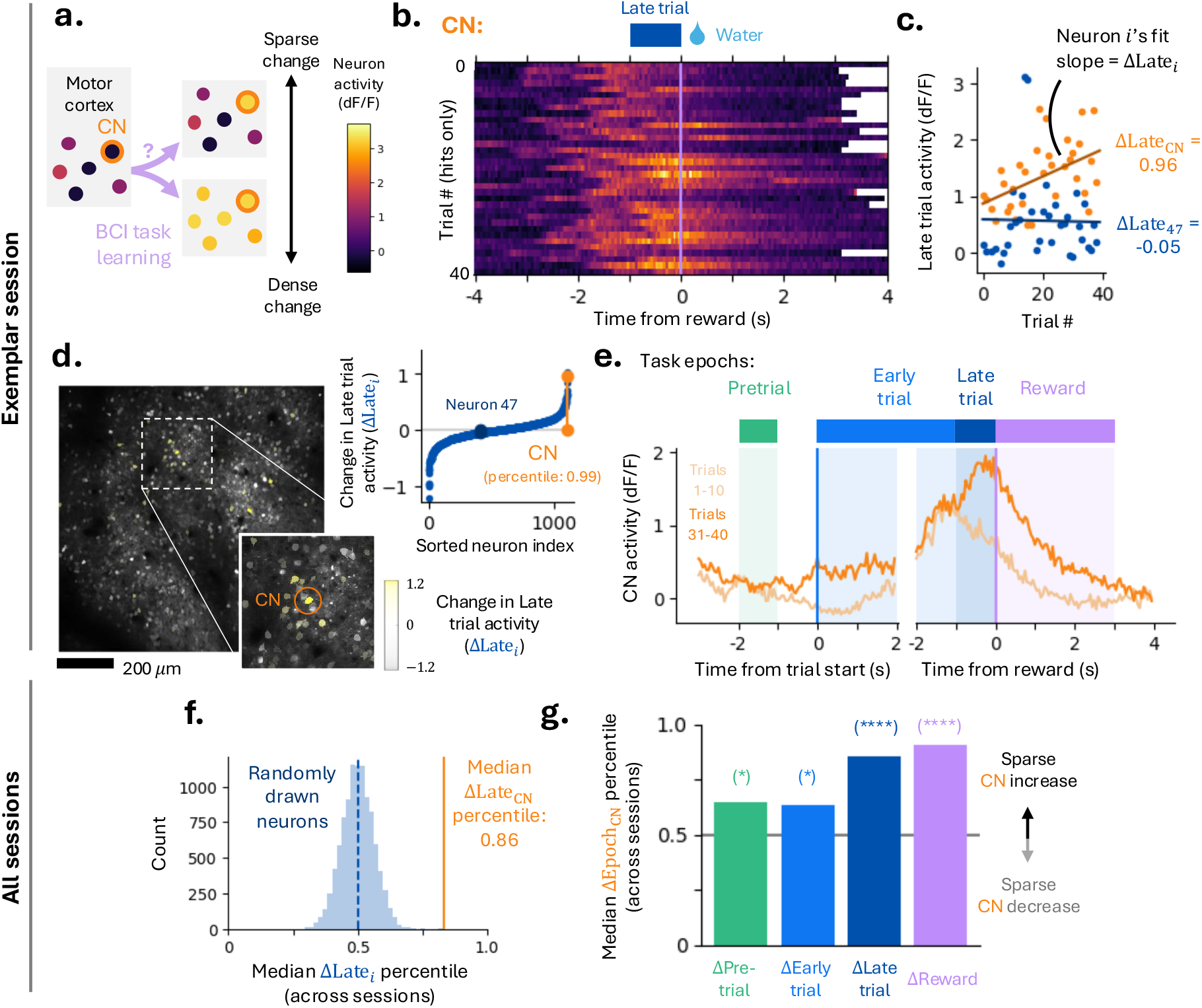
Changes in neural activity are sparse. **[a-e]** Exemplar session. **(a)** Schematic illustrating two possible strategies for BCI learning: sparse modulation of the CN versus dense population activity changes. **(b)** CN activity aligned to reward delivery, plotted versus trial #. Miss trials are omitted. The late trial epoch is defined as the 1 s interval preceding reward. Same color scale as (a). **(c)** Mean late trial activity of the CN and an exemplar neuron across trials. Slope of linear fits were used to quantify change in late trial activity (ΔLate_*i*_ for neuron *i*). **(d)** Left: FoV calcium fluorescence with neuron ROIs shaded by change in late trial activity (ΔLate_*i*_); inset highlights CN. Right: ΔLate_*i*_, with CN and exemplar neuron indicated. ΔLate_*i*_ of the CN is in the 99th percentile. **(e)** Top: definition of task epochs relative to trial start and reward delivery. Bottom: Mean CN activity across trials 1-10 and 31-40, aligned to trial start (left) and reward delivery (right), illustrating changes across epochs. **[f-g]** Aggregate data across sessions. **(f)** Distribution of median ΔLate_*i*_ percentiles across all sessions for randomly sampled neurons from each session (10^4^ samples). Orange line indicates median CN percentile. **(g)** CN median change in task epoch activity (i.e. ΔPre_*i*_, ΔEarly_*i*_, ΔLate_*i*_, ΔRew_*i*_) percentile across sessions for all four task epochs. Significance was estimated via bootstrap (10^4^ draws; from left to right: *p* = 0.011, 0.017, < 10^−4^, < 10^−4^).

Neuronal activity was aligned to salient cues during trials, e.g. time of reward (exemplar CN, Fig. 2b). We first focus on the 1 s interval immediately preceding reward (‘late trial’ epoch, Fig. 2b), during which reward-port speed is high (Fig. 1h). Quantifying trial-by-trial changes in each neuron’s late trial activity using the slope of a linear fit, we found that the CN exhibited large within-session increases in late trial activity (Fig. 2c). Across the FoV, the majority of neurons showed little learning-related change in this epoch (Fig. 2c-d), indicating that learning-related modulation is sparse. Sparse late trial activity changes were consistent across sessions, and tended to be larger for neurons spatially proximal to the CN (Fig. S2), consistent with the columnar connectivity in MC (Finkelstein et al. 2025).

To determine whether CN-specific changes were restricted to the late trial epoch, we divided the trial into four epochs defined by task events (Fig. 2e): (1) pretrial, the inter trial period terminated by trial start; (2) early trial, following the auditory cue but preceding late trial; (3) late trial, the final 1 s before reward; and (4) reward, immediately after reward delivery (Methods). Comparing median late trial changes across mice and sessions to a null distribution, the CN consistently exhibited significantly larger increases than randomly sampled neurons (median percentile: 0.86, 1st–3rd quartile: 0.15-0.98; bootstrap *p* < 10^−4^; Fig. 2f). This selectivity persisted when the null distribution was restricted to neurons that would have been viable CN candidates at session onset (low trial-start tuning and elevated activity; Fig. S2). Extending this analysis to other task epochs, the CN showed a pronounced increase in reward-epoch activity, reflecting lingering late trial activity (Fig. 2g). In contrast, CN modulation during pretrial and early trial epochs was minimal (Figs. 2g, S2). These results indicate that BCI learning engages a small subset of neurons, including the CN, with selective modulation confined to behaviorally relevant epochs.

### Signatures of upstream and motor cortex plasticity in RNNs

Are changes in CN activity driven by local plasticity within the MC network, or by plasticity involving upstream regions providing input to MC (Fig. 3a)? To address this, we built recurrent neural network (RNN) models to test whether sparse activity changes are more consistent with either upstream or MC plasticity. The CN is selected from the RNN hidden layer. This layer represents MC, and its recurrent connections model the local circuit. The input population provided task-relevant signals, corresponding to higher-order cortical and subcortical inputs.

**Figure 3:**
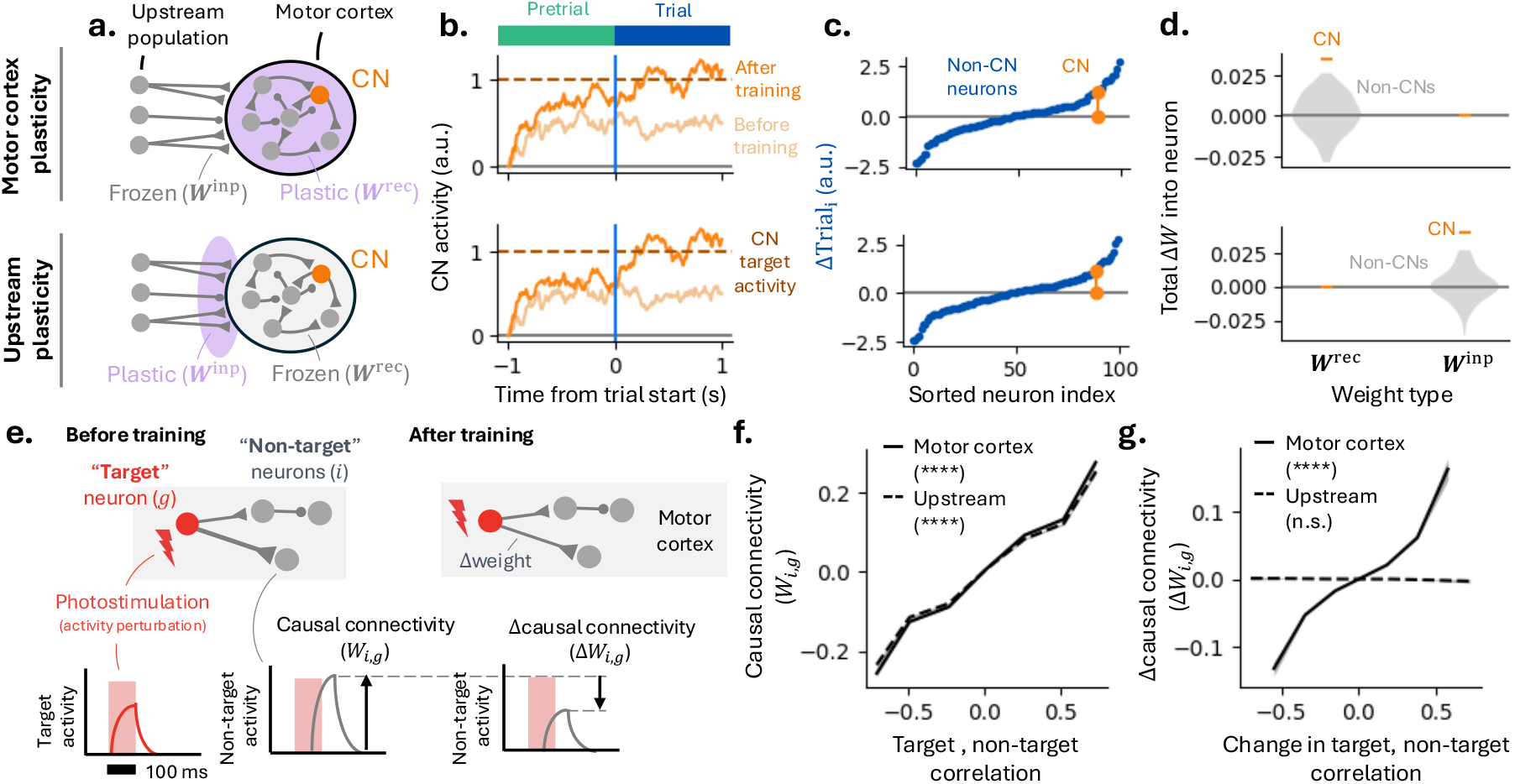
Signatures of upstream and motor cortex plasticity in recurrent neural networks (RNNs). **[a-d]** ‘Motor cortex’ (MC) plasticity model (top) versus ‘upstream’ plasticity model (bottom). **(a)** RNN models that differ in which weights are modified via training. The MC plasticity model posits that changes in CN activity are caused by synaptic plasticity (purple) between neurons in the local MC surrounding the CN. In the upstream plasticity model, changes occur at synapses from distal inputs onto MC. **(b)** CN activity increases over training in both MC and upstream models. **(c)** Both models exhibit sparse, CN-targeted activity changes (example sessions). **(d)** Weight changes, quantified as the sum of synaptic modifications onto each postsynaptic neuron, for CN and non-CN neurons. **(e)** Schematic of weight measurement via simulated perturbations (analogous to 2-photon photostimulation). Left and right panels show network before and after training. ‘Target’ neurons (indexed by *g*) are stimulated, producing responses in ‘non-target’ neurons (indexed by *i*), quantified as ‘causal connectivity’ (*W*_*i,g*_). Weight changes are inferred by comparing *W*_*i,g*_ before and after training (‘Δcausal connectivity’, Δ*W*_*i,g*_). **(f)** Across both MC (solid line) and upstream (dashed) models, pairwise activity correlations predict causal connectivity (*p* < 10^−4^, positive Wald test). **(g)** Only the MC plasticity model (solid line) exhibits a relationship between learning-related *changes* in correlations and *changes* in causal connectivity (*p* < 10^−4^, positive Wald test).

The RNN was tasked with increasing CN activity during the trial. Plasticity was restricted to different synaptic populations to model upstream versus MC learning. In the MC plasticity model, only recurrent synapses were plastic, whereas in the upstream model, only input-to-MC synapses were modified (Fig. 3a). These scenarios represent extremes along a continuum in which both pathways could contribute to learning.

When trained with backpropagation through time (BPTT), both models learned the BCI task within tens of trials by increasing CN activity before and after trial onset (Fig. 3b), and both exhibited sparse learning (Fig. 3c). In the upstream model, synaptic changes were concentrated on neurons projecting to the CN, whereas in the MC plasticity model, modifications occurred primarily in recurrent synapses converging onto the CN (Fig. 3d). In both cases, additional changes to non-CN neurons influenced CN activity indirectly via recurrent dynamics.

Thus, RNN models indicate that rapid learning and sparse activity changes are consistent with either MC or upstream plasticity. We next used our models to determine whether targeted photostimulation (Chettih & Harvey 2019; Marshel et al. 2019; Russell et al. 2024; Daie et al. 2021; Carrillo-Reid et al. 2019) could be used to distinguish between upstream and MC plasticity (Fig. 3e). Targeted photostimulation perturbs neuronal activity to probe how neurons influence others in the same circuit, through monosynaptic and polysynaptic pathways. In the RNN, perturbations applied to hidden-layer neurons modeled MC photostimulation, and resulting changes in surrounding activity defined a map of ‘causal connectivity’ (Fig. 3e).

We examined how causal connectivity relates to activity correlations between neuron pairs during task performance. In both models, neurons with higher activity correlations exhibited stronger causal connectivity, consistent with recurrent MC connections shaping shared activity patterns (Ko et al. 2011; Chettih & Harvey 2019; Daie et al. 2021; Finkelstein et al. 2025) (Fig. 3f).

Next, we tested whether learning-related *changes* in activity correlations predicted *changes* in causal connectivity. If learning modifies recurrent connectivity within the recorded MC population, positive correlations are expected because both correlation structure and causal connectivity depend on connection strength between MC neurons. On the other hand, upstream plasticity can alter local correlations, but because it does not modify recurrent MC connections these changes are not predictive of the changes in causal connectivity that arise from modifying upstream inputs. Consistent with this, in the MC plasticity model, changes in correlation strength predicted changes in causal connectivity, whereas this relationship was absent in the upstream model (Fig. 3g). Models incorporating plasticity in *both* upstream and MC pathways produced intermediate outcomes, depending on the relative magnitude of synaptic changes in each layer (Fig. S3). An alternative upstream model, in which plasticity was restricted to recurrent connections within the upstream network, yielded similar results (Fig. S3).

### Mapping causal connectivity across days using two-photon photostimulation and imaging

We interleaved BCI learning with repeated mapping of causal connectivity. At the end of each BCI session, we used holographic two-photon photostimulation and imaging to map causal connectivity among L2/3 MC neurons within the field-of-view (Fig. 4a). Holographic photostimulation pulses (100 ms duration) were delivered every 600 ms to random groups of ten neurons, with 100 distinct groups each repeated ~20 times (Methods). In addition to directly photostimulated neurons, other neurons in the FoV exhibited changes in fluorescence amplitude following photostimulation of the targeted group (Fig. 4b).

**Figure 4:**
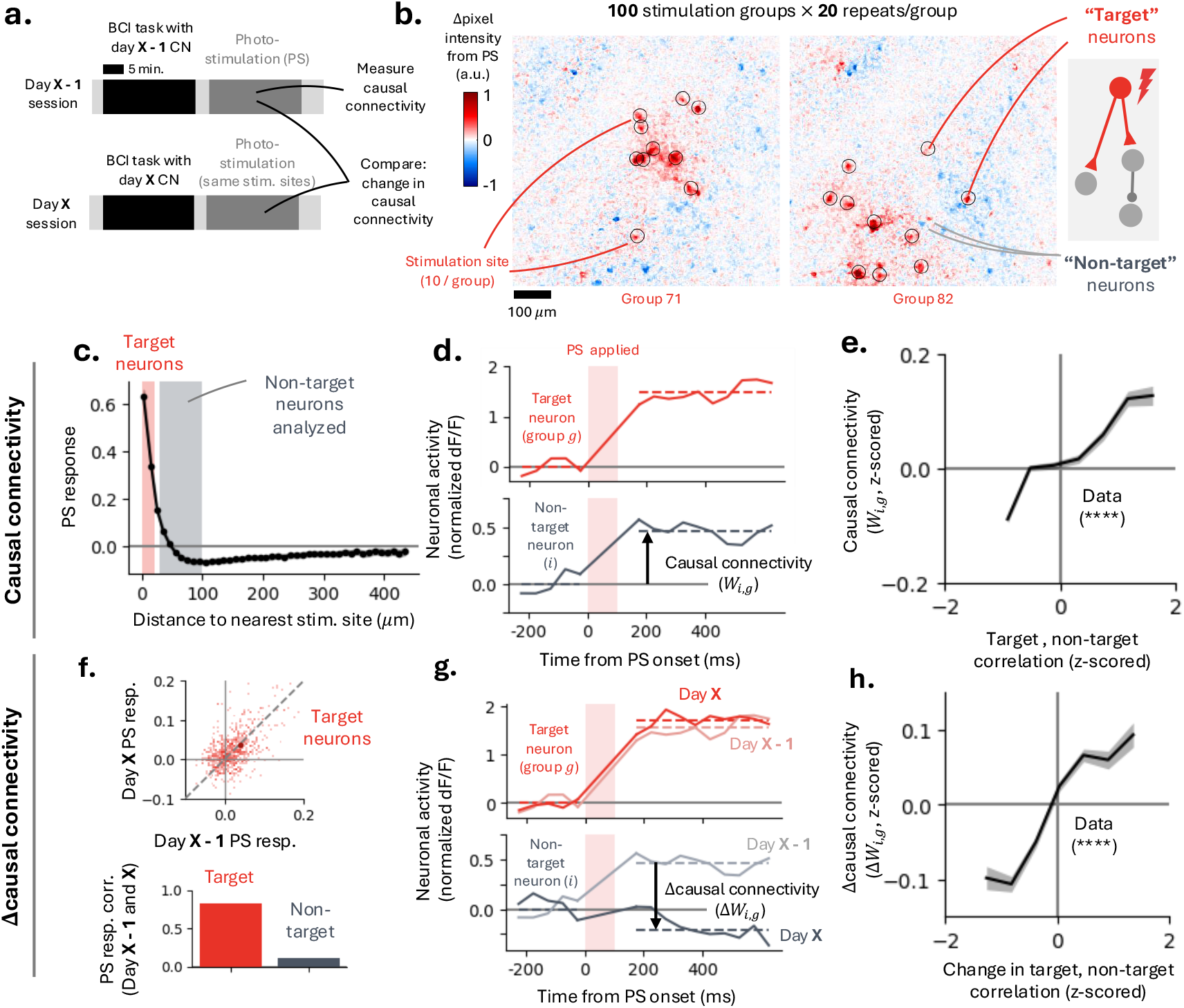
Mapping causal connectivity across days reveals local MC plasticity. **(a)** Experimental design: photostimulation (PS) mapping was performed following BCI training on each day. **(b)** Example photostimulation experiment showing the mean change in neuronal activity before and after PS for two stimulation groups. Each mapping session comprised 100 distinct stimulation groups with 10 stimulation sites, each repeated approximately 20 times. **[c-e]** Measurement of causal connectivity. **(c)** PS-evoked activity change as a function distance from the nearest stimulation site. Neurons within 20 µm of a site were classified as ‘target neurons’. Analyses of ‘non-target’ neurons were restricted to neurons located 30-100 µm from the nearest stimulation site. **(d)** Mean activity change of target neuron (top) and a nearby non-target neuron (bottom) following photostimulation of group *g*. ‘Causal connectivity’ (*W*_*i,g*_) quantifies PS-evoked activity change of non-target neuron *i*. **(e)** Causal connectivity of non-target neurons as a function of the activity correlation between target and non-target neurons during the BCI task (binned data, mean ± s.e.m.; positive Wald test, *p* < 10^−4^). **[f-h]** Changes in causal connectivity. **(f)** Top: scatter plot of target neuron PS responses measured on consecutive days. Bottom: across-day correlations in PS responses for target and non-target neurons. **(g)** Mean activity change of a target neuron (top) and a nearby non-target neuron (bottom) evoked by photostimulation of group *g* on two consecutive days. ‘Δcausal connectivity’ (Δ*W*_*i,g*_) quantifies across-day change in the response of non-target neuron *i*. **(h)** Δcausal connectivity of non-target neurons as a function of changes in activity correlations between target and non-target neurons during the BCI task (binned data, mean ± s.e.m.; positive Wald test, *p* < 10^−4^).

Photostimulation-evoked activity (PS responses) decreased with distance from the photostimulation site: on average, neurons within ~60 µm exhibited positive (excitatory) responses, whereas neurons beyond this distance showed weaker and often negative (inhibitory) responses (Fig. 4c). To separate direct from indirect effects of photostimulation, neurons within 20 µm of a stimulation site were classified as ‘target neurons’ (potentially directly stimulated), while neurons 30–100 µm away were classified as ‘non-target neurons’ (Daie et al. 2021). The ‘causal connectivity’ (*W*_*i,g*_) of each non-target neuron (*i*) to a photostimulation group (*g*) was defined as the change in its activity following photostimulation, normalized by its repeat-to-repeat variability (Fig. 4d, Methods). All subsequent analyses focused on non-target neurons to isolate causal connectivity from direct photostimulation effects.

Neuronal correlations during the BCI task predicted causal connectivity measured immediately afterward (18 sessions, 6 mice, 1.3 × 10^5^ target-non-target pairs). Non-target neurons exhibiting stronger positive (negative) correlations with target neurons during behavior showed correspondingly stronger excitatory (inhibitory) causal connectivity (Fig. 4e; positive Wald test, *p* < 10^−4^, ρ = 9.25 × 10^−2^). These results indicate that activity correlations in MC during task performance reflect the underlying structure of MC connectivity, consistent with previous findings (Chettih & Harvey 2019; Daie et al. 2021; Finkelstein et al. 2025).

### Changes in causal connectivity reveal local MC plasticity

To probe local MC plasticity, we compared causal connectivity maps obtained on consecutive days. Photostimulation was performed at the end of each day to measure MC connectivity before and after BCI learning on the later day (Fig. 4a). The same stimulation groups were repeated across days, enabling longitudinal measurements of causal connectivity between the same sets of neurons (12 session pairs, 6 mice, 8.2 × 10^4^ target-non-target pairs). Across days, PS responses of target neurons were strongly correlated (Fig. 4f), consistent with stable direct photostimulation effects. In contrast, PS responses of non-target neurons, whose responses reflect synaptic connectivity, exhibited substantially weaker day-to-day correlations.

For each non-target neuron (*i*) and stimulation of group *g*, we defined across-day changes in causal connectivity (‘Δcausal connectivity’, Δ*W*_*i,g*_) as the difference between the two days’ PS-evoked responses, normalized by the pooled repeat variance across both days (Fig. 4g, Methods). Changes in activity correlations between target and non-target neurons during the BCI task predicted corresponding changes in causal connectivity (Fig. 4h; positive Wald test, *p* < 10^−4^, ρ = 6.12 × 10^−2^). This relationship indicates that neurons whose activity became more (less) correlated during learning subsequently exhibited stronger (weaker) causal connections. Thus, our photostimulation experiments provide direct evidence for changes in causal connectivity in MC induced by BCI learning (Fig. 3g).

### Diverse population task tuning across BCI task epochs predicts causal connectivity

We next asked if causal connectivity is organized around task-relevant tuning.

Aligning neuronal activity to trial start and reward onset revealed heterogeneous temporal modulation across the neural population. Individual neurons were modulated at distinct times during the trial, such that the population collectively tiled the entire trial (Fig. 5a-b). Unlike the CN, whose activity directly determines BCI control, other MC neurons influence task performance indirectly through the local MC circuit, including the CN.

**Figure 5:**
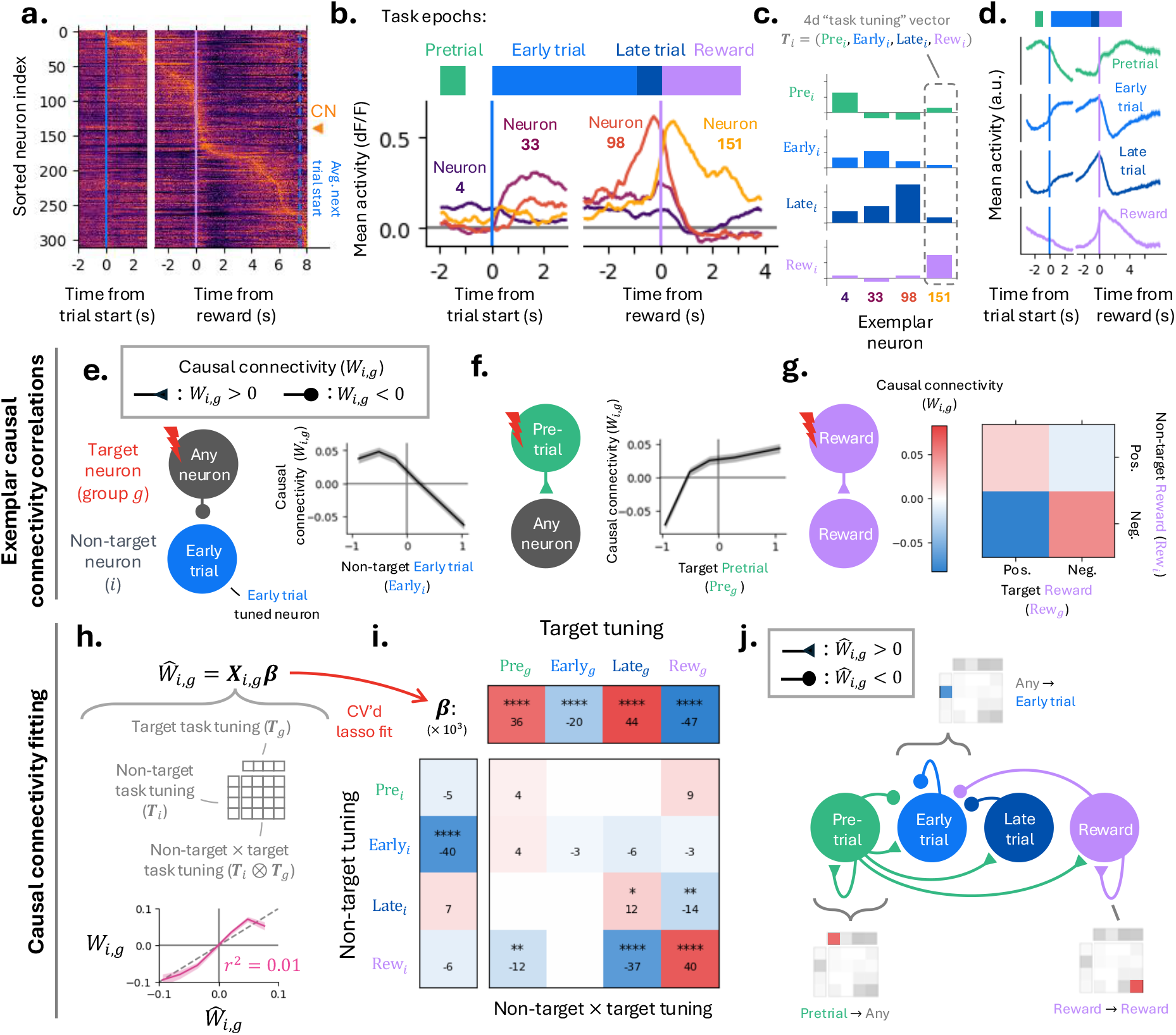
Task tuning during BCI training predicts causal connectivity. **(a)** Neuronal activity aligned to trial start and reward delivery aligned activity for neurons in field-of-view, sorted by time of peak activity. The activity of each neuron is *z*-scored for visualization. **(b)** Example neurons showing trial-averaged activity with tuning to distinct task epochs (pretrial, early trial, late trial, and reward). Colored bars indicate task epoch definitions throughout figure. **(c)** The activity of each neuron is summarized by a four-dimensional ‘task tuning’ vector, **T**_*i*_ = (Pre_*i*_, Early_*i*_, Late_*i*_, Rew_*i*_), defined as the mean activity of each neuron within each task epoch. **(d)** Population-averaged trial activity, computed by weighting the activity of each neuron by its corresponding task tuning component (e.g. Pre_*i*_ for the pretrial epoch; mean ± s.e.m. across sessions). **[e-g]** Exemplars illustrating functional specificity of causal connectivity (binned data, mean ± s.e.m.). **(e)** Causal connectivity varies with the task tuning of the non-target neuron (example shown for early trial tuning). **(f)** Causal connectivity depends on the task tuning of the target neurons (example shown for pretrial tuning). **(g)** Causal connectivity is stronger between target and non-target neurons with similar task tuning (example shown for reward tunings). **[h-j]** Regression analysis of task tuning and causal connectivity. **(h)** Causal connectivity is fit using multivariate linear regression (MLR) with predictors derived from the task tuning vectors of non-target neurons (**T**_*i*_), target groups (**T**_*g*_), and their interactions (**T**_*i*_ ⊗ **T**_*g*_), including exemplar features illustrated in [e-g]. Cross-validated lasso regression selects a sparse model; prediction performance on held-out data is shown below. **(i)** Regression coefficients from the MLR, indicating which task-tuning features significantly predict causal connectivity (*t*-test significance shown). **(j)** Schematic summarizing the connectivity patterns implied by salient regression coefficients in (i). Notation denotes effective causal connectivity influence, which may reflect multi-synaptic interactions.

We summarized the task tuning of each neuron by stacking their mean activity across the four task epochs introduced earlier (pretrial, early trial, late trial, and reward) into a four-dimensional ‘task tuning’ vector (**T**_*i*_; Fig. 5c). To visualize the population-wide temporal structure implied by these tunings, we computed epoch-weighted averages of activity across neurons, using the corresponding task-tuning component of each neuron (e.g. Pre_*i*_ for pretrial; Fig. 5d).

Population-level pretrial tuning was elevated following reward and persisted through inter-trial delay until trial start, resembling preparatory activity observed in movement tasks (Churchland et al. 2006; Shenoy et al. 2011; Churchland et al. 2012). early trial and late trial tuning peaked after trial start and before reward, with low activity in the inter-trial interval. reward tuning was transiently elevated following reward delivery. Thus, population dynamics transitions between distinct, epoch-specific states over the course of the trial (Fig. S5).

We next examined how task tuning relates to causal connectivity. Specifically, does causal connectivity (*W*_*i,g*_) depend on the task tuning of its non-target neuron (*i*) and/or target group (*g*)?

We first provide three illustrative examples showing a relationship between task tuning and causal connectivity. First, non-target neurons with stronger early trial tuning exhibited smaller (more inhibitory) photostimulation-evoked response changes following target stimulation, corresponding to weaker causal connectivity (Fig. 5e). Second, stronger pretrial tuning in target neurons predicted larger causal connectivity to non-target neurons (Fig. 5f). Third, for target and non-target neurons with similar reward tuning—both positively or both negatively tuned— causal connectivity was stronger than when their tunings differed, indicating preferential connectivity between similarly tuned neurons (Fig. 5g).

These observations motivated a systematic analysis of the relationship between causal connectivity and (1) the non-target task tuning vector, (2) the target task tuning vector, and/or (3) the similarity between the two task tuning vectors. To assess these dependencies, we constructed a multivariate linear regression (MLR) model relating causal connectivity to task tuning. Predictors included the non-target task tuning vector (**T**_*i*_), the target-group task tuning vector (**T**_*g*_, defined as a weighted sum of the task tuning vectors of target neurons in group *g*), and all pairwise interactions between their components (i.e., the outer product **T**_*i*_ ⊗ **T**_*g*_), yielding 4 + 4 + 4 × 4 = 24 candidate predictors (Fig. 5h). Lasso regularization was applied to promote sparse fits, with the regularization parameter selected by cross-validation. The resulting regression predicted held-out causal connectivity values (Fig. 5h), indicating that a compact subset of task tuning features captures key aspects of MC connectivity.

The regression coefficients quantify how task tuning relates to causal connectivity (Fig. 5i). Positive coefficients indicate that stronger tuning is associated with more excitatory causal connectivity, whereas negative coefficients are associated with more inhibitory connections. The example correlations observed in the raw data (Fig. 5e-f), non-target early trial suppression, target pretrial excitation, and like-to-like reward connectivity also emerged as significant predictors in the regression (Figs. 5j). Together, these results show that population task tuning across task epochs predicts structured patterns of causal connectivity.

### Pretrial-tuned neurons dominate BCI-related plasticity

We next extended this framework to predict *changes* in causal connectivity across days. Because BCI control is driven by the CN and earlier we observed increases in its late trial activity (quantified by a Δlate trial> 0), we hypothesized that learning would preferentially strengthen connections from neurons already active during the late trial epoch onto neurons that increase their late trial activity, which includes the CN (Fig. 6a). Such changes would be consistent with like-to-like connectivity, whereby strengthened connections from neurons tuned to a given feature induce increases in that tuning in their targets.

**Figure 6:**
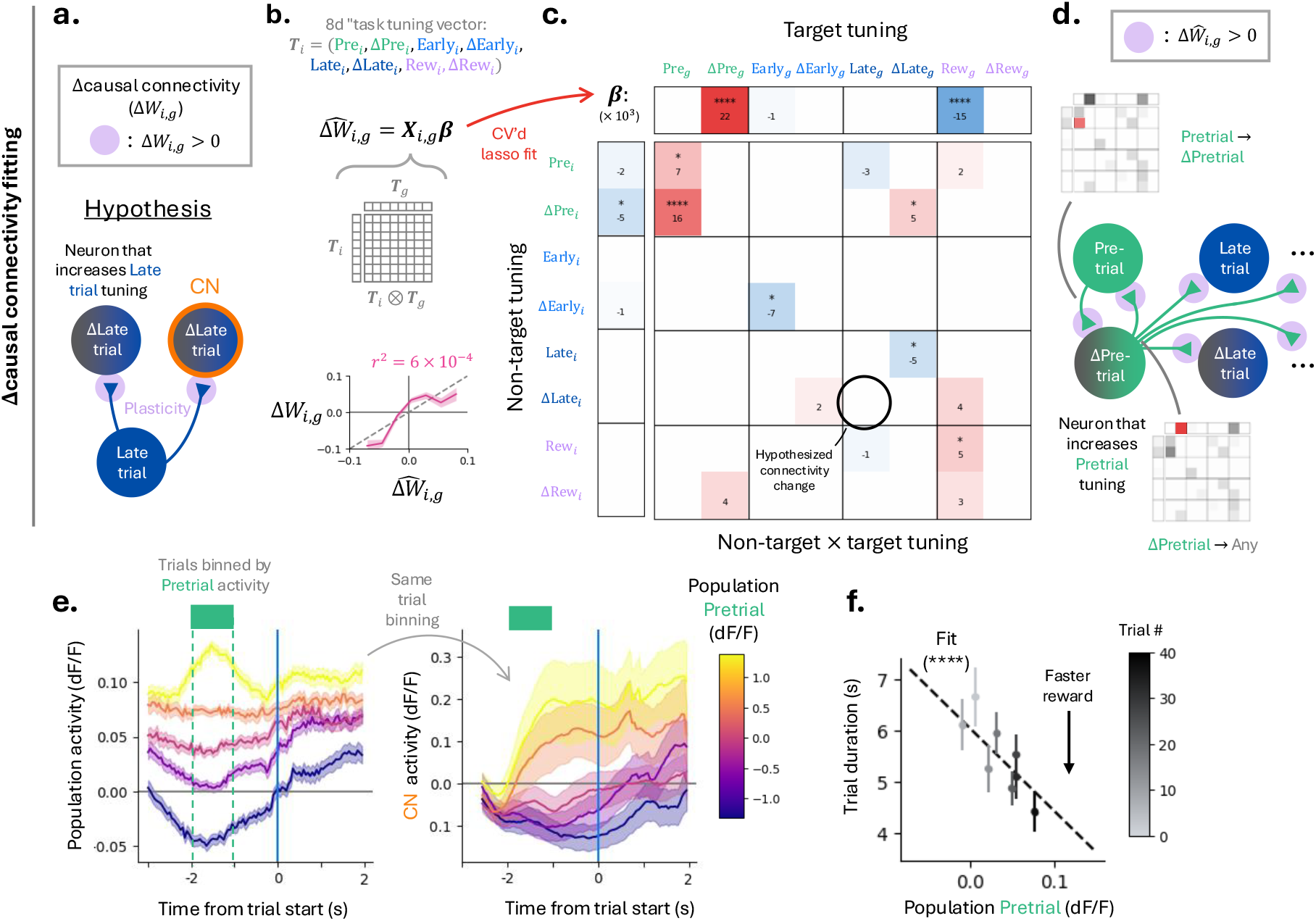
Changes in causal connectivity are enriched in neurons participating in preparatory activity. **(a)** Hypothesized circuit-level mechanism for BCI learning: enhanced late trial tuning, which includes the CN, could arise from strengthened connectivity from neurons already tuned to the late trial epoch. **[b-d]** Regression analysis of Δcausal connectivity. **(b)** Δcausal connectivity is fit using multivariate linear regression (MLR) with predictors derived from both target and non-target task tuning vectors. The fit parallels the regression used for causal connectivity in Fig. 5h, with additional regressors capturing learning-related changes in task tuning. Cross-validated lasso regularization selects a sparse model; prediction performance on held-out data is shown below. **(c)** Regression coefficients from the MLR, indicating which neurons task tuning features and tuning changes predict Δcausal connectivity. **(d)** Schematic summarizing connectivity changes implied by salient regression coefficients in (c). **[e-f]** Functional role of pretrial activity during the BCI task. **(e)** Activity aligned to trial start, sorted according to the amplitude of pretrial activity (CN omitted). Mean population activity (left) and CN activity (right) are shown, demonstrating that pretrial population activity predicts subsequent population and CN activity during trial. **(f)** Trial duration as a function of population pretrial activity, showing that higher pretrial activity correlates with improved performance. Points are colored by trial #, illustrating that pretrial activity increases over the session alongside behavioral improvement (*p* < 10^−4^, Wald test).

To identify which learning-related changes in neuronal tuning predict changes in causal connectivity, we expanded the task tuning vector to include changes in task tuning. Specifically, we appended epoch-specific tuning changes (ΔPre_*i*_, ΔEarly_*i*_, ΔLate_*i*_, and ΔRew_*i*_), previously used to quantify learning-related activity changes (Fig. 2), to the original four-dimensional task-tuning vector. This yielded an eight-dimensional task tuning vector that captures both a neuron’s baseline task tuning and how that tuning shifts during BCI learning.

We used these extended task tuning vectors in an MLR model to predict Δcausal connectivity (Δ*W*_*i,g*_). As before, regressors included the non-target task tuning vector, the target-group task tuning vector, and all pairwise interactions between their components (i.e., the outer product). With eight-dimensional task tuning, this resulted in 80 candidate predictors. Sparsity was again enforced using cross-validated lasso regularization. The resulting model predicted held-out Δcausal connectivity values (Fig. 6b), indicating that task tuning and learning-related changes forecast structured changes in MC connectivity.

The fitted regression coefficients reveal which neuronal properties (task tuning, or change in task tuning) are most predictive of connectivity changes (Fig. 6c). Target neurons that increased their pretrial tuning tended to exhibit larger positive Δcausal connectivity to non-target neurons (Fig. 6d), consistent with the earlier finding that strong pretrial tuning in a target neuron is associated with stronger excitatory connections. In contrast to our initial hypothesis, we did not detect preferential strengthening of connectivity from late trial tuned neurons onto neurons that increased their late trial tuning (Fig. 6c, circled). Instead, an analogous pattern emerged for the pretrial epoch: target neurons with strong pretrial tuning preferentially strengthened their causal connections to non-target neurons that acquired or increased pretrial tuning (Fig. 6c, arrow).

Importantly, refitting the causal connectivity MLR with the additional tuning-change predictors yielded coefficients largely consistent with those obtained using baseline task tuning vectors alone, indicating that changes in activity are generally not strongly predictive of static causal connectivity measurements (Fig. S5). However, these tuning changes are important for explaining changes in causal connectivity across learning.

Together, these results demonstrate that MC plasticity during BCI learning preferentially strengthens connections among neurons participating in pretrial activity. Activity preceding trial start may therefore play a privileged role in coordinating subsequent task-related tuning and learning-related reconfiguration of MC circuits (Perich et al. 2018; Sun et al. 2022).

### Pretrial activity reflects preparatory dynamics predictive of trial state and performance improvement

Motivated by the dominant role of pretrial activity in predicting changes in causal connectivity, we next asked if pretrial activity exhibits hallmarks of preparatory dynamics-namely, whether it predicts upcoming neural and behavioral states during the BCI task (Churchland et al. 2006; Shenoy et al. 2011; Churchland et al. 2012).

Following reward delivery on the preceding trial, population activity remained elevated until trial start, with additional modulation time-locked to the cessation of reward consumption (Fig. S6). This sustained pretrial activity suggests that the pretrial population state constrains the initial conditions from which control-related dynamics unfold. Consistent with this interpretation, population activity during the pretrial epoch predicted both subsequent population-wide activity and CN activity following trial start (Fig. 6e, S6).

pretrial activity was also predictive of behavioral performance: trials preceded by higher pretrial activity were shorter in duration. Moreover, pretrial activity increased across trials, paralleling improvements in task performance (Fig. 6f). These results indicate that pretrial activity exhibits preparatory dynamics that set initial conditions for trial evolution and support learning-related improvements in BCI performance. Together, these results support a picture in which pretrial neurons form a preparatory state that drives upcoming trial activity. This is supported by connectivity measurements showing that pretrial neurons provide net excitatory drive to the rest of the network, and by the observation that learning-related plasticity is concentrated within the Δpretrial population.

### A model circuit explaining BCI learning

pretrial activity predicted both learning-related changes in causal connectivity and subsequent improvements in task performance. Plasticity acting on neurons participating in preparatory dynamics contribute to BCI control. We asked whether the connectivity changes observed experimentally could, on their own, enable BCI learning.

We constructed a minimal network model of the transition from pretrial to late trial activity, incorporating weight changes that reproduce key features of the measured causal connectivity and Δcausal connectivity (Methods; Figs. 5–6). For simplicity, the model transitions directly from the pretrial to the late trial epoch, effectively collapsing EARLY and late trial responses as would occur in short BCI trials where the former becomes negligible.

Before learning, the network is wired to reproduce a subset of the causal connectivity motifs observed in the data (Figs. 7, S7), including the widespread excitation coming from the pretrial population driving a downstream late trial population (Fig. 7a) (Fig. 5j). Consistent with experimental data, the model CN initially exhibits weak late trial tuning, with activity remaining below the reward port movement threshold (Fig. 7b).

**Figure 7:**
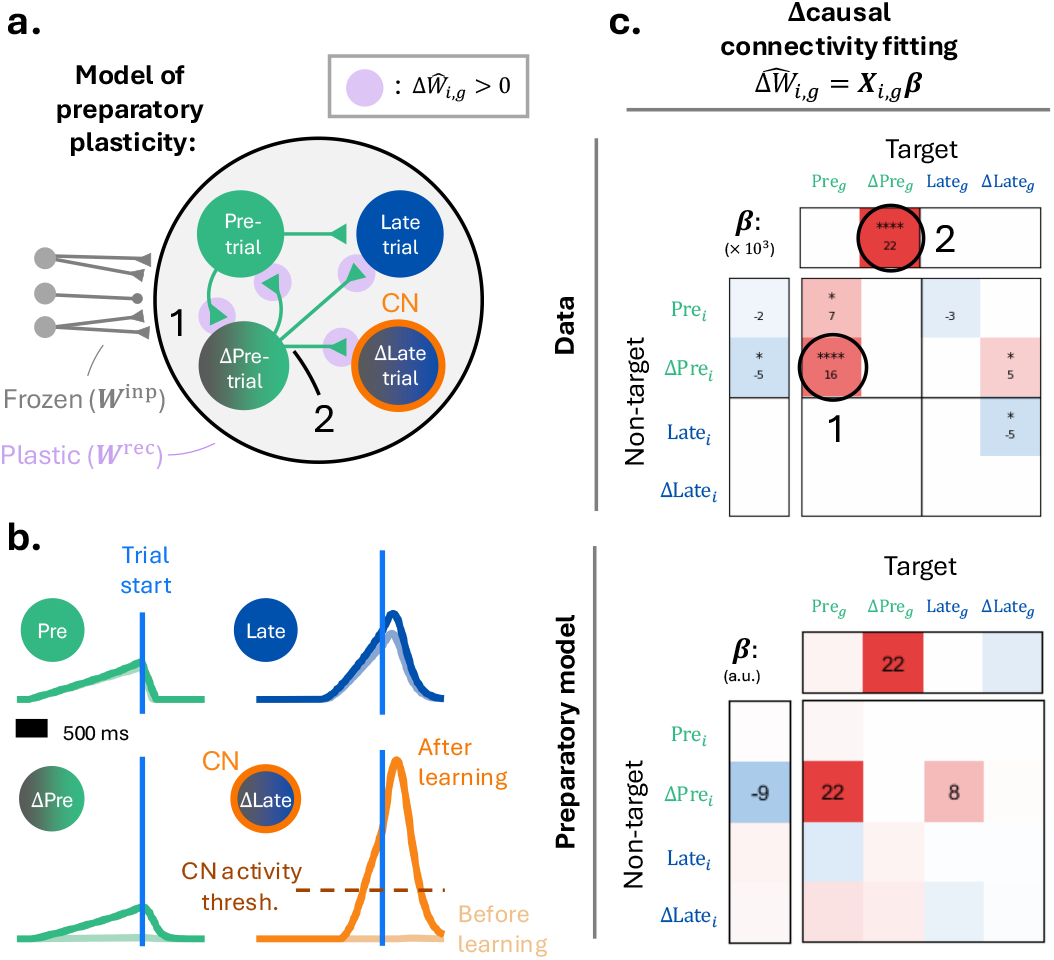
A preparatory model of BCI learning. **(a)** Schematic diagram of the preparatory network (Methods). Magenta circles indicate connections that are strengthened during learning to match characteristics seen in data, labeled by two of the prominent changes seen in the data. **(b)** Trial activity of pretrial, Δpretrial, late trial, and Δlate trial (e.g. the CN) tuned neurons shown in (a) before (light) and after (dark) learning. **(c)** Top: Subset of MLR coefficients fitting Δcausal connectivity, shown in Fig. 6b, to be matched by the model. Labels correspond to those in (a). Bottom: MLR coefficients fitting model Δcausal connectivity (colored by size of coefficient, a.u.).

We then introduced learning-related connectivity changes motivated directly by the Δcausal connectivity fits. Strengthening inputs from existing pretrial-tuned neurons onto a subset of untuned neurons causes these neurons to acquire elevated pretrial activity, forming a Δpretrial population (Fig. 7a, label ‘1’). This manipulation reproduces a prominent feature of the empirical Δcausal connectivity fit: increased connections from pretrial neurons onto Δpretrial neurons (Fig. 7c, label ‘1’). Next, we added a broadcasted increase in excitation from these Δpretrial neurons to the rest of the network, reproducing additional features of the measured Δcausal connectivity (Figs. 7a, label ‘2’). This additional preparatory drive increases late trial tuning across the population, including in the CN (Fig. 7b, bottom right). As a consequence, CN activity crosses the movement threshold, leading to increased reward port movements and successful BCI control. An alternative connectivity model in which Δpretrial preferentially connect back onto the original pretrial population yields similar results (Fig. S7).

These conceptual models suggest that BCI learning does not require recruiting arbitrary new downstream targets. Instead, learning can proceed by selectively strengthening interaction within an existing preparatory network, expanding the Δpretrial population and amplifying its influence on subsequent trial dynamics. This strategy preserves the low-dimensional organization of motor cortical activity while enabling effective activation of the CN and improved task performance.

## Discussion

BCI learning in motor cortex produced rapid (Fig. 1) and sparse (Fig. 2) changes in neural activity, accompanied by changes in causal connectivity (Figs. 4, 6). The observed relationship between connectivity changes and neural correlations was consistent with local MC plasticity, but not with purely upstream plasticity (Figs. 3-4). Although local MC plasticity may be sufficient to drive the rapid learning observed during BCI, it does not exclude contributions from upstream areas to learning-related changes in MC activity. Plasticity was concentrated among neurons that were active during the pretrial epoch, which exhibited preparatory activity (Fig. 6). These preparatory neurons, which establish the network’s initial state before trial start, appear to play a central role in supporting rapid BCI learning through local circuit plasticity (Fig. 7).

The dynamics of MC activity during BCI learning have features that resemble MC activity during motor control. For example, MC activity occupies distinct preparatory and movement-related states (Kaufman et al. 2014; Elsayed et al. 2016), with transitions between these states typically triggered by a ‘go’ cue in laboratory tasks. The preparatory state serves as the initial condition for the ensuing motor command (Churchland et al. 2012). During BCI learning, pretrial activity similarly predicts subsequent neural activity and behavioral performance. Models of motor control predict strong drive from neurons active in the preparatory mode onto neurons that drive movement (Hennequin et al. 2014). Consistent with this prediction, connectivity mapping in the intact brain reveals both correlational and causal influences from pretrial neurons onto the broader local population, including neurons engaged during the trial. Together, these findings suggest that preparatory activity provides a widespread drive that helps establish the initial conditions for task execution.

Several studies have suggested that motor learning and BCI learning are associated with a reorganization of preparatory activity. For example, preparatory activity in motor cortex changes in response to a remapping between sensory inputs and motor outputs (Perich et al. 2018; Sun et al. 2022). Modeling work implies that such shifts in preparatory activity could arise from changes in the pattern of inputs to the preparatory network (Menéndez et al. 2025). In contrast, our data and modeling support an alternative mechanism: plasticity *within* MC is sufficient to drive the learning-related shift in preparatory activity. Our model network explains activity changes based on the connectivity changes observed in the data. In this network, the preparatory neurons integrate and amplify external pre-trial inputs. Modifying the connections to and from the preparatory neurons increases subsequent trial activity in the condition neuron.

These findings reveal that rapid, functionally meaningful reconfiguration of local circuitry can occur on the timescale of behavioral improvements (a few behavioral trials). Similarly, recent work has shown that motor learning can rapidly generate new neural activity patterns in MC, beyond the flexibility expected from reassociation on a stable manifold (Sun et al. 2022). Such rapid reorganization is consistent with known synaptic mechanisms operating on short timescales (Markram et al. 1997; Froemke et al. 2010; Sjöström et al. 2001). The slower emergence and stabilization of new synapses likely reflects a subsequent consolidation phase rather than the initiation of learning (Fu et al. 2012; Xu et al. 2009).

Multiple theoretical frameworks have proposed how performance feedback shapes synaptic plasticity during learning (Legenstein et al. 2010; Lillicrap et al. 2020; Lillicrap et al. 2016; Williams 1992). These models differ in their biological implementation, speed of learning, and precision of credit assignment. Discriminating among these models has been difficult due to the scarcity of direct measurements of circuit-level plasticity during learning. Our combined BCI and photostimulation-based connection mapping helps bridge this gap by linking behavioral learning to measured changes in local MC connectivity.

Concentrating plasticity within the preparatory sub-network may support rapid learning by restricting changes to a low-dimensional subspace of the circuit. Small synaptic adjustments within this sub-network can produce large, coordinated changes in task-related activity, while leaving most of the network unchanged. This organization may allow the system to adapt quickly without overwriting existing motor representations, preserving the global structure of motor cortical dynamics while enabling flexible behavioral reconfiguration.

How is plasticity biased toward the preparatory neuron population? One possibility is that this pattern reflects targeted credit assignment, in which neurons participating in preparatory activity are preferentially modified because changes in their synapses exert a disproportionate influence on subsequent network activity. Another possibility is that plasticity is selectively gated in these neurons, for example through cell-type–specific properties, such as selective access for neuromodulatory input.

Taken together, these results demonstrate that rapid learning can arise from the selective reorganization of local MC circuitry. By integrating all-optical connectivity mapping with a precisely controlled learning paradigm, we link synaptic-level plasticity to emergent population dynamics and behavioral adaptation. This framework offers a mechanistic bridge between theories of network-level learning and cellular plasticity, and establishes a foundation for uncovering how local circuit modifications enable flexible, goal-directed behavior.

## Methods

### Experiment

#### Mice

Data are from 10 mice (age at the beginning of experiments ~70-150 days) and a total of 78 behavioral sessions. Half of the mice were CamK2a-tTA (JAX, catalog no. 007004) × Ai94 (TITL-GCaMP6s; JAX, catalog no. 024104) × slc17a7 IRES Cre (JAX, catalog no. 023527). The other half of mice were Camk2a-tTA x tetO-GCaMP6s (JAX, catalog no. 024742).

#### Surgical procedures

All procedures were in accordance with protocols approved by the Allen Institute and Janelia Research Campus Institutional Animal Care and Use Committees. Cranial window surgeries were performed as described previously (https://dx.doi.org/10.17504/protocols.io.bqstmwen). Prior to surgery, mice were injected with Dexamethasone (intramuscular) and Ceftriaxone (subcutaneous). During surgery, mice were anesthetized with 1-2% isoflurane. A 3 mm craniotomy was drilled above forepaw primary motor cortex (1 mm anterior and 2 mm lateral from Bregma). 50 nL AAV2/2 camKII-KV2.1-ChrimsonR-FusionRed (Addgene, 10^12^ GC/ml; plasmid catalog no. 102771) was injected in a 3 × 3 grid pattern centered at the center of the craniotomy with 600 micron spacing, 300 microns deep from the surface of the brain. The craniotomy was then covered by 3 layers of 3 mm circular glass. The window and a custom headplate were attached to the skull using cyanoacrylate glue and metabond (Parkell). Mice were started on water restriction three weeks after surgery and were given at least 5 days to acclimate to restriction before behavioral experiments.

#### Two-photon calcium imaging and photostimulation

Two-photon imaging and photostimulation were performed on a commercial microscope (Bergamo, Thorlabs) with a customized holographic light path through a 16x Nikon objective (Fig. S1). The photostimulation path consisted of a 1035 nm pulsed laser (Monaco, Coherent). A spatial light modulator (SLM; 1920 × 1152; Meadowlark) was mounted on an optical breadboard on top of the microscope. The SLM was imaged onto a pair of galvanometer mirrors using 2 inch lenses in a 4f configuration (Packer et al. 2015). Imaging was performed with a 920 nm laser (Chameleon, Coherent) was modulated by a Pockels cell (Conoptics) and its position was controlled with a resonant-linear galvanometer pair. The imaging and photostimulation beams were combined before the objective using a 1000 nm long pass filter. Imaging and photostimulation were both controlled using Scanimage (MBF Biosciences).

Fields of view were either 800 × 800 or 1000 × 1000 µm depending on the spatial extent of ChrimsonR expression. Images were acquired at 20 Hz (800 × 800 pixels). Prior to experiments, a stack of imaging planes was acquired centered on the FoV (21 planes, 2 µm spacing, for each plane 100 frames were acquired and averaged). These image stacks were referenced in real-time during experiments to correct for brain motion.

Two-photon imaging data were analyzed using Suite2p (Pachitariu et al. 2017). Fluorescence activity was computed as the mean intensity within each ROI and expressed as Δ*F*/*F*_0_, with *F*_0_ defined as the median fluorescence across the session. We use *h*_*i,t*_ and *ĥ*_*i,t*_ to respectively denote the Δ*F*/*F* and raw fluorescence activity of neuron *i* at time *t*.

#### Videography and part tracking

Behavioral videos were acquired using two cameras (FLIR) positioned beneath and to the side of the mouse. Videos were acquired at 400 Hz using the pySpincapture GUI. Movies were analyzed using a Deeplabcut model trained on 400 total frames from multiple mice.

#### Brain-computer interface

ROIs were defined in Scanimage using the Integration module. Average fluorescence intensity of all pixels within ROIs was tracked in real-time. The activity of the CN (*ĥ*_CN_; average intensity of all pixels within the ROI) converted to an analog voltage (*V*_*m*_, 0-3.3 V) to control the frequency (*f*_step_) of steps (each 685 µm) towards the mouse of the motorized reward port according to the equations

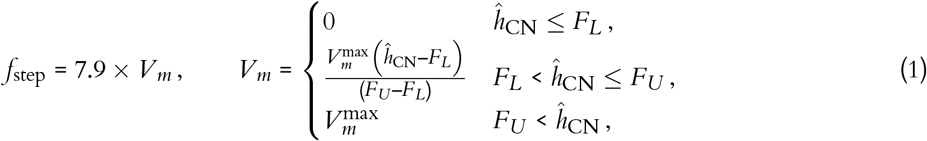

where *F*_*U*_ and *F*_*L*_ are the upper and lower threshold which were calculated as the maximum and median fluorescence values during the 1-5 minutes long spontaneous period immediately preceding the BCI behavioral epoch (Fig. S1). The Scanimage motion correction module was used to adjust the position of the CN ROI in real-time. This online motion-correction introduced an additional delay between CN activity and reward port movements of approximately 500 ms (mean 530 ms; standard deviation 270 ms).

#### Trial structure

At the start of each trial the reward port begins 7 mm away from the mouth of the mouse (‘start position’). Trial initiation was indicated by an auditory cue (Fig. S1). Water reward could be released from the reward port with tongue contact only after the reward port reached the ‘reward position’ (6 mm from the start position) within the trial duration (10 seconds). The speed of the reward port was controlled by CN activity. The time of tongue contact is defined as the reward time. If the reward position was not reached within 10 seconds, a new trial was initiated. To initiate a new trial after a rewarded trial, the activity of the CN had to return below *F*_*L*_ for at least 200 ms, which initiated a delay of 2 seconds before the next trial start. No additional stimulus cues were given to indicate these transitions. At trial start the reward port moved from the reward position to the start position in approximately 300 ms. Once the port reached the start position, the next trial was initiated. Mice performed 105 trials per session (median, 1st-3rd quartile: 95-136) lasting 22.0 minutes (median, 1st-3rd quartile: 17.3-25.5).

#### CN selection

We selected conditioned neurons (CNs) based on two criteria: activity modulation throughout the session and trial start tuning. Activity modulation was calculated as the fraction of time spent above the standard deviation of activity (⟨*h*_*i,t*_ > σ_*i*_ ⟩_*t*_). The median CN was active during 9.2% of the recording session prior to becoming the CN (median, 6.1%-12.1%, 75% CI) which represented the 97.6th percentile across all recorded neurons (median, 90.1th - 99.3th percentile, 75% CI). Trial start tuning (i.e. the mean activity between trial start and trial end, minus the mean activity in the epoch (−2 s, −1 s) before trial start) of the CNs prior to conditioning was 0.05 (median, 0.01—0.21, 75% CI) which represented the 38th percentile (median, 7.6th–87.7th percentile, 75% CI) across recorded neurons. Weaker trial start tuning tended to reduced initial reward rates and thus increase task difficulty.

#### Photostimulation

To map local circuit connectivity *in vivo*, we combined holographic 2-photon photostimulation and imaging. We simulated both single and multi-neuron photostimulation to determine which most accurately reconstructed causal connectivity and found that reconstruction speed increased with the number of neurons stimulated simultaneously, saturating at approximately ten neurons per group (Fig. S4). Based on these modeling results, we photostimulated groups of 10 neurons per photostimulus in experiments.

Photostimuli were applied sequentially every 600 ms. Experiments consisted of 100 different photostimulation groups. Each photostimulation group consisted of 10 photostimulation sites centered on neurons selected at random with a spatial constraint requiring the maximum separation between any pair of neurons to be less than 400 µm. This spatial constraint was implemented due to physical limitations of the SLM (Yang et al. 2016). The photostimulation beam was scanned over the target cell bodies in a spiral pattern (Rickgauer & Tank 2009) lasting 10 ms and repeated 10 times (100 ms total duration). The same photostimulation group is photostimulated several times during a photostimulation sessions over distinct ‘repeats’ (median: 20, 1st-3rd quartile:17-23). In addition to neurons whose centers are the photostimulation sites, neurons within 20 µm of the photostimulation beam showed significant direct responses to photostimulation (Chettih & Harvey 2019; Daie et al. 2021; Finkelstein et al. 2025). We therefore treat all neurons within 20 µm of a stimulation site as ‘target’ neurons. Neurons that are more than 30 µm away from the nearest photostimulation site are considered to potentially indirectly respond to photostimulation (Daie et al. 2021; Finkelstein et al. 2025). When assessing changes in causal connectivity, we photostimulate the same 100 groups each session after BCI learning (12 session pairs, 6 mice).

### Analysis

All trial-dependent analyses were restricted to the first 40 trials of each session, as performance improvements were typically observed early in a session, with increasing hit rates and decreasing trial durations across sessions and mice (Figs. 1f, S1).

We note that the increases in CN activity that reduced the trial duration were also associated with longer inter-trial intervals (mean increase: 1.15 s, 0.48–1.82, 95% CI), reflecting elevated post-reward CN activity that delayed decay below *F*_*L*_ and the next trial start. Despite this slowdown, mice exhibited robust improvements in hit rate, reward rate, and time-to-reward.

#### Task epochs, tuning vectors, and changes in activity

We define four task epochs: pretrial, early trial, late trial, and reward. These distinct epochs were motivated by salient trial stimulus modulation seen in the data. The pretrial epoch was defined to be between 2 and 1 seconds before trial start in order to capture activity after the low CN activity threshold had been met and before the reward port returned to start position, indicating the initiation of a new trial. Since the trial initiation conditions differed for hit and miss trials (Fig. S1), pretrial response was only computed following hit trials. The division of a trial into early trial and late trial was motivated by the fact that a significant portion of reward port movement happens at the very end of the trial (Fig. 1h), and so we aimed to separate modulations from the auditory trial start cue and significant reward port movement. Motivated by the greater-than average reward port movement period (Fig. 1h), the late trial epoch occurs 1 to 0 seconds before reward delivery, and thus only occurs on rewarded trials. The reward epoch occurs from 0 to 3 seconds after reward delivery (and thus only on hit trials) to capture initial reward delivery modulation as well as subsequent reward consumption, which lasts several seconds (Figs. 1b, 6e).

To quantify activity modulation in each task epoch and how they change over a session, neuronal activity is aligned to trial start and reward times (Fig. 2b). The mean activity of each neuron over the task epoch is computed for each trial, e.g. 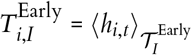 is the early trial activity of neuron *i* on trial *I* and *I* = 1, …, 40 indexes the trials of a session. Here, ⟨·⟩𝒯denotes a temporal average over times 𝒯and 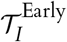 are the early trial times of trial *I*.

We define the change in task epoch activity of neuron *i* (e.g. ΔEarly_*i*_) as the slope of the (ordinary least square) linear regression fit of 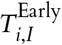 as a function of *I* (Fig. 2c; similarly defined for other task epochs). Likewise, mean task epoch activity is computed for each neuron over all trials, e.g. 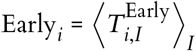 (Fig. 5c). Thus the mean activity modulation of neuron *i* to all four task epochs can be summarized by the 4 numbers (Pre_*i*_, Early_*i*_, Late_*i*_, Rew_*i*_) and its change in activity over the session by the 4 numbers (ΔPre_*i*_, ΔEarly_*i*_, ΔLate_*i*_, ΔRew_*i*_). These quantities are combined into the four- and eight-dimensional ‘task tuning’ vectors used in the main text. As a control, task tuning vectors defined on only half of trials (odd/even) are predictive of tuning on the other half (Fig. S5).

To quantify how changes in CN activity across sessions compared to the rest of the population, we bootstrapped a distribution by randomly drawing a single neuron *i* from each session and then computing the median ΔLate_*i*_ across all sessions (Fig. 2f). This was then compared to the median ΔLate_*i*=CN_ across sessions. Importantly, since CNs are not chosen randomly across all neurons (see CN selection above), we also computed bootstrapped distribution when drawing *i* from neurons that would be considered candidate CNs at the start of the session (Fig. S2a). To do so, for a given CN, we looked at both its activity magnitude and its tuning around trial starts in the session directly prior to the one where it was chosen to be the CN. We then created the bootstrap distribution by only drawing from the subset of neurons that had the most similar activity magnitude and tuning around trial start to the chosen CN (5% of all neurons). This process was repeated for the other task epochs in Figs. 2, S2). When this analysis was repeated for the CN from the previous day, no significant changes were observed (Fig. S2).

#### Causal connectivity and Δcausal connectivity

Let the index *g* = 1, …, 100 refer to the photostimulation groups; *a*_*g*_ be the photostimulation (PS) repeat for photostimulation group *g* (*a*_*g*_ = 1, …, 20 on average, but can vary slightly between groups); and α be the session index. We drop the α superscript when distinction between sessions is not needed. Here, we begin by quantifying the response to photostimulation of all neurons within the FoV (which we refer to as ‘PS response’) before formally defining the causal connectivity and Δcausal connectivity that only applies to a subset of neurons and is analyzed throughout the main text. This distinction will be important for controlling for changes in stimulation strength in target neurons.

To quantify the response of all neurons to photostimulation, we compute the difference between the activity of each neuron before and after photostimulation application, averaged across time. We define the ‘repeat response’for neuron *i* to repeat *a*_*g*_ of photostimulation group *g* during session α to be

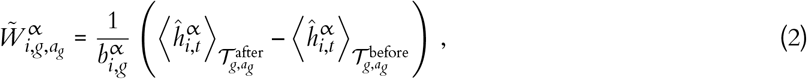

where 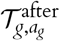 are the set of times 300 ms after the photostimulation and 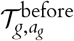 the set of times 200 ms before the *a*_*g*_ repeat of photostimulation group *g* (Fig. S4). We exclude responses for the 100 ms during the photostimulation because response measurement is contaminated by the photostimulation laser. Here, 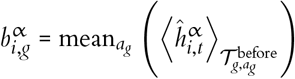 is the baseline activity for each neuron before photostimulation, averaged across repeats. 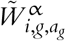 is normalized by the average pre-stimulation response to account for fluorescence differences from day to day. Additionally, if a neuron was a target neuron in the previous photostimulation application, we do not compute its repeat response in the following photostimulation application, since we found they often still had elevated activity. We define the ‘PS response’ of each neuron to be its mean across repeat responses normalized by its standard deviation across repeats,

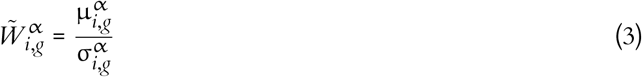

where 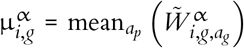 and 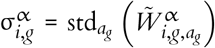 denote the average and standard deviation across the photostimulation events (Fig. S4). This normalization ensured that photostimulation responses that were highly variable were down-weighted/less significant compared to those neurons that had a more regular/more significant responses (Fig. S4). Similar normalizations have been used previously in the literature (Chettih & Harvey 2019).

The change in PS response (‘ΔPS response’) to stimulation of group *g* is defined as

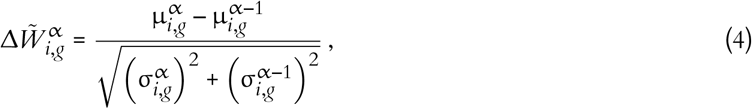

where we now normalize the response by the squared sum of variances, a standard way of propagating errors across differences. Computing ΔPS response requires photostimulation of the same photostimulation sites on subsequent days, which reduces our set of possible sessions to 12 pairs of sessions across 6 mice. PS and ΔPS responses are only defined for neuron/group combinations that meet a minimum repeat number threshold, which we set to be 10 throughout this work.

Connection strength and probability has a strong distance dependence, with the strongest excitatory connections being within 100 microns (Song et al. 2005; Daie et al. 2021; Perin et al. 2011; Campagnola et al. 2022; Finkelstein et al. 2025). These connections and their changes are the target measurement of the photostimulation in this work. We define two subsets of neurons for each PS group, *g*. Let represent the 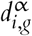 minimum distance of neuron *i* to the photostimulation beam in group *g* during session α. We respectively define the set of ‘target’ and ‘non-target’ neurons for group *g* to be

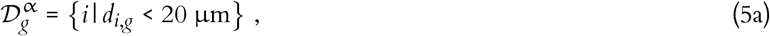

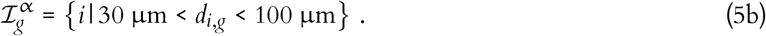

When we analyze pairs of sessions to compute changes in photostimulation response, we use the same target and non-target neuron masks across both sessions. Since the PS sites for a given PS group are the same neurons on both days (see above), the vast majority of neurons will be considered target/non-target neurons on both days from the definitions above. For a neuron to be considered target/non-target, we require it to meet the criterion of Eq. (5) across both sessions. Neurons that do not meet these criterion are omitted from analysis. In practice, the neuronal distances from photostimulation sites we use in our definition of target/non-target are significantly correlated across days, with only 4.9% and 2.8% of target and non-target neurons changing definition across days (and thus being omitted), respectively (median; ranges: 0.8-17.5% and 0.6-7.3%, respectively).

PS and ΔPS responses can be computed for any neuron in the FoV for a given stimulation group. In the main text, we restrict the analysis of PS responses to non-target neurons and refer to this subset of PS responses and ΔPS responses as ‘causal connectivity’ (*W*_*i,g*_) and ‘Δcausal connectivity’ (Δ*W*_*i,g*_), respectively:

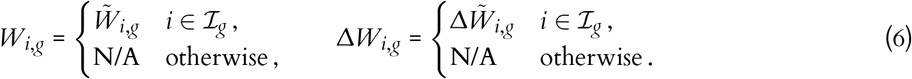

#### Causal connectivity fitting

Throughout the main text, we analyze various measures to explain causal connectivity or Δcausal connectivity in non-target neurons. We are interested in understanding how the task tuning of (1) the non-target neuron, (2) the photostimulation group, and/or (3) the similarity of the non-target neuron and photostimulation group affect the causal connectivity.

Since we photostimulate groups of neurons rather than single neurons, properties of the photostimulation groups will be weighted sums over all target neurons within the group.^1^ For example,

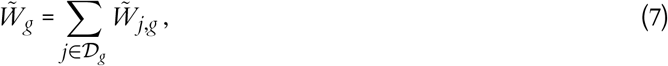

represents the total PS response for all target neurons in group *g* when group *g* is photostimulated. We find how much the target neurons respond to photostimulation is strongly correlated with how much non-target neurons respond to the same stimulation (Fig. S4).

We first sought to understand if correlations between target and non-target neurons during the BCI task were explanatory of the photostimulation responses of the non-target neurons. To compute the correlation of a given non-target neuron with the target neurons in photostimulation group *g*, we take a weighted sum over the pairwise correlation between the non-target neuron and all target neurons for that group,

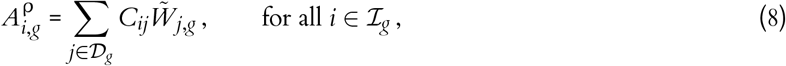

where *C*_*ij*_ = ρ (*h*_*i*_, *h*_*j*_) is the Pearson correlation between the activity of neurons *i* and *j* during the BCI session. Although we consider all neurons within 20 µm of the photostimulation beam to be target neurons, a large fraction of these neurons do not exhibit a PS response, potentially due to a differences in expression or location along the hologram, and thus their stimulation may have variable influence on non-target neurons. To account for this variability in target neuron response, we weighted correlations between target and non-target neurons by the responses of the target neurons in Eq. (8). This weighting ensures that we excluded target neurons with weak or zero responses to photostimulation from the target to non-target neuron correlation.

Given the influence of the summed target response has on the non-target response, we were concerned that metrics of the form of Eq. (8) could reflect a spurious association arising from the known dependence of *W*_*i,g*_ on the total target response, rather than an independent effect of Eq. (8). For example, if mean_*j*_ (*C*_*ij*_) ≠ 0, *W*_*i,g*_ might be well explained by Eq. (8) simply because it is well explained by Eq. (7). To verify this is not the case, we fit *W*_*i,g*_ to both these quantities simultaneously using multivariate linear regression (MLR), and see if there remains significant evidence for correlation influencing the non-target responses after the total target response magnitude has been regressed out.We indeed find that correlations explain causal connectivity even when we control for this potential additional source of influence (Fig. 4e). We adopt this control procedure, i.e. regressing out contributions from target PS response size, throughout this work.

In order to understand causal connectivity beyond correlations, we considered how they depend on target and non-target task tuning vectors. Generalizing the form of Eq. (8), we developed a class of regressors of the form,

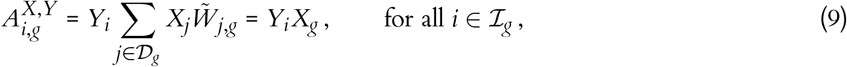

where *Y*_*i*_ and *X*_*j*_ may be elements of the non-target and target task tuning vectors, respectively. Here we have defined 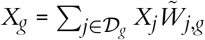 which represents the PS response-weighted group task tuning vector. For example, for a regressor to probe for the dependence shown in Fig. 5e, we would take *Y*_*i*_ to be the early trial tuning (Early_*i*_) of the non-target neuron and *X*_*g*_ = 1, since it has no dependence on the target neuron properties. Similarly, for the dependence of Fig. 5g, we would take *Y*_*i*_ to be the reward tuning (Rew_*i*_) of the non-target neuron and *X*_*j*_ to be the reward tuning (Rew_*j*_) of the target neuron.

The MLR fit shown in Fig. 5h involves fitting *W*_*i,g*_ with regressors of the form Eq. (9) with all possible combinations of *X*_*i*_ and *Y*_*g*_ being either 1, Pre_*i*_, Early_*i*_, Late_*i*_, or Rew_*i*_ (equivalently with *g* subscripts, with the *Y*_*i*_ = 1 and *X*_*g*_ = 1 case redundant with the intercept). Including the control regressor, Eq. (7), yields a total of 25 regressors. Cross validation fits to determine optimal L1 regression coefficients are done over 10 folds.

#### Δcausal connectivity fitting

Similar to the fitting of causal connectivity, we next aimed to understand if task tuning could explain the Δcausal connectivity (Δ*W*_*i,g*_). Like causal connectivity fitting, we found the changes in causal connectivity were influenced by the change in target stimulation between days (Fig. S4). Thus we defined the change in photostimulation response of target neurons in group *g* as

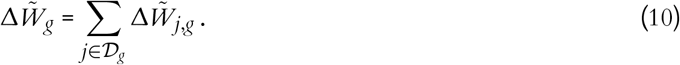

Like the control of the causal connectivity fitting, we find this regressor to be significantly predictive of Δ*W*_*i,g*_ (Fig. S4). Thus we include this control in all fits henceforth, effectively regressing out this contribution to the Δcausal connectivity.

Given the significant relationship between photostimulation response of non-target neurons and their correlation with the target neurons, we were motivated to determine if changes in correlation were predictive of changes in the non-target photostimulation response. We define the change in correlation between non-target neuron *i* and photostimulation group *g* to be

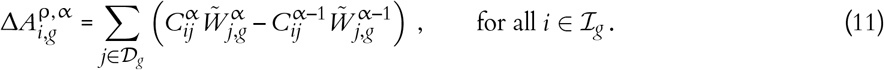

Fitting Δ*W*_*i,g*_ using MLR with this change in correlation along with our control regressor, Eq. (10), we indeed find a significant relationship between changes in correlation and causal connectivity (Fig. 4h).

For Fig. 6b, the regressors take the same form as Eq. (9) but now also append ΔPre_*i*_, ΔEarly_*i*_, ΔLate_*i*_, ΔRew_*i*_ (i.e. a now eight-dimensional task tuning vector). All combinations of *Y*_*i*_ and *X*_*g*_, along with the control regressor, Eq. (10), yields a total of 81 regressors (again omitting *Y*_*i*_ = 1 and *X*_*g*_ = 1) for fitting Δ*W*_*i,g*_. All other MLR procedures are conducted in a manner analogous to the causal connectivity fitting setup.

To assess raw correlations in both the causal connectivity and Δcausal connectivity to regressors of the form of Eq. (9), we also performed MLR fits containing the control regressor and only one of the 80 regressors at a time (Fig. S6).

### Models

To represent learning within and upstream of the motor cortex, we use a firing-rate recurrent neural networks (RNNs) trained using supervised backpropagation through time (BPTT).

#### Network

We use Vanilla RNNs to model the cortical circuit, with the hidden layer (*n* = 100 neurons) representing neurons in the MC and the input layer (*d* = 10 neurons) representing upstream inputs onto the MC. Hidden layer neurons are indexed with *i, j* = 1, …, *n* and input layers neurons with *I, J* = 1, …, *d*. The hidden activity from one time step to the next is given by

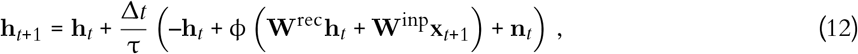

where **h**_*t*_ and **x**_*t*_ are the hidden and input activity of the RNN, respectively. Here, Δ*t* = 0.01 s is the time step, τ = 0.1 s the network time constant, ϕ (·) = tanh (·) the non-linearity. **n**_*t*_ ~ 0.01 × 𝒩 (0, 1) represents additional input noise injected into the hidden layer.

The BCI activity is the activity of a single neuron, via a one-hot mask, 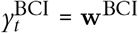 **h**_*t*_ where **w**^BCI^ is a one-hot vector that selects out the CN (see below for CN selection details).

All weights in the network are drawn from normal distributions. Specifically, 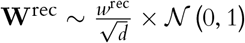 where *w*^rec^ = 0.5 was chosen so that the network did not have particularly long-lived modes during photostimulation, which was observed empirically. Likewise, 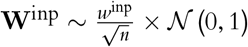 where *w*^inp^ = 0.1. We found that *w*^inp^ < *w*^rec^ in order to observe the positive relationship between correlation and causal connectivity seen in the data (Fig. 4e).

#### BCI Task

We train our model networks on individual trials representative of the BCI trials, where performance is measured by how close the CN can reach the target activity. Each trial is divided into only two periods, representative of the pretrial and late trial period of the experiment. Distinct constant inputs are provided during the pretrial and late trial periods.

The target activity 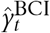 is defined to be high during the late trial, and is calibrated based on the spontaneous activity of the CN during untrained trials (see below). No target activity is given during the pretrial period. All hidden activity is reset to 0 at the start of each trial.

The CN is determined based on hidden activity during a 10 trial activity stabilization period where no weight adjustment occurs. The CN is chosen from the set of neurons that have above median pretrial and late trial activity, specifically the neuron whose difference in pretrial and late trial is smallest. These conditions mimic the requirements from experiment that the CN must be a neuron that is both active and has small tuning around trial start. We then define 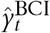 to be the 95th percentile of CN activity during the 10 trial stabilization period, also mimicking the experimental definition from spontaneous activity.

#### Training

Upstream and local MC plasticity only train **W**^inp^ and **W**^rec^, respectively. Training begins and ends with a 10 trial stabilization periods where no weight adjustment occurs. These periods are used to analyze mean network activity under the noisy inputs present during training. The models are trained using supervised BPTT, which tries to minimize the MSE loss of 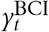 with respect to the target BCI activity 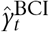. Models were trained one trial at a time (batch size was equal to 1).

Learning rates were scanned over to yield training rates comparable to one another and what was observed in experiment (Fig. S3). Whether training **W**^inp^ or **W**^rec^, the RNN models were able to increase CN activity to close to the target activity within 30 trials, mimicking the rapid learning speed of the mice. However, since the network only needs to change the activity of the CN to minimize loss and BPTT has direct knowledge of what neuron is the CN through **w**^BCI^, we found that activity and weight changes were far more confined to the CN than what was observed in experiment (Fig. S3). To encourage more widespread changes, we used feedback misalignment so that backpropagation credit was fed through a layer **w**^back^ ≠ **w**^BCI^ (Lillicrap et al. 2016). Specifically we take 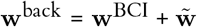 where 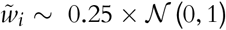 for all *i* that are not the CN. This ensures **w**^back^ · **w**^BCI^ = 1 so training still occurs (Lillicrap et al. 2016). With this modification, weight changes were far more widespread but training speed could still be calibrated to match experiment.

#### Photostimulation

Although we have direct access to connection strengths in models, photostimulation is applied before and after learning to mimic experiment. Photostimulation is applied to hidden neurons individually instead of in groups, so the number of groups is equal to the number of hidden neurons. A perturbation is applied by clamping the activity of a given neuron to a fixed value, *h*^pert^, for the duration of the photostimulation. *h*^pert^ is fixed across all neurons and photostimulation repeats. The photostimulation response, 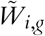, is defined as the change in the activity of neuron *i* before and during the photostimulation application (*g* now indexes individual neurons, but we keep this notation for consistency). Noise is injected through the input layer during photostimulation, otherwise no additional input representing stimuli is provided. 5 photostimulation repeats were performed for each stimulation group (neuron). Since we have no spatial embedding and can precisely and reliably stimulate neurons, we consider all neurons that are not directly perturbed to be non-target neurons, i.e. 𝒟= *g* and ℐ_*g*_ = {*i* | *i* ≠ *g*}.

#### Analysis

Correlations between hidden layer neurons, *C*_*ij*_ are computed during the 10 trial stabilization periods before and after training. Since the direct photostimulation response is the same for all directly stimulated neurons, our control regressors 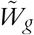 and 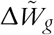 are not needed for the models nor are the target neuron PS response weights used in regressors of the form of Eq. (9).

#### Alternative models

Among additional models we consider in the supplement (Fig. S3), we train a network with both **W**^inp^ and **W**^rec^ plastic. We adjust the learning rate such that weight changes between the two were comparable.

We also train a network that can adjust the excitability of each neuron, where Eq. (12) is replaced with:

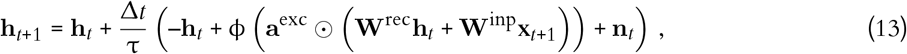

with **a**^exc^ the excitability of each neuron and ⊙the element-wise product. **a**^exc^ = **1** at initialization, and all other weights are left fixed. We find this training to be significantly noisier than the other setups, so omit networks with no decrease in loss (17/100 initializations rejected).

We also considered a model where the upstream inputs were generated via an RNN:

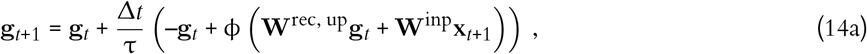

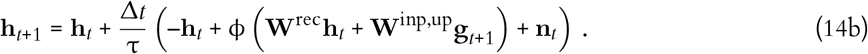

with **g**_*t*_ ∈ ℝ^20^ the upstream hidden units that serve as input into the RNN representing the motor cortex. In this case, only **W**^rec, up^ was left plastic. We found that this training was also noisy and rejected initialization that did not decrease their loss (9/100 initializations rejected).

Finally, we also consider a network with additional recurrent connections from the hidden layer back to the input layer, potentially representing cortical-subcortical loops that may be present from the motor cortex back to upstream inputs, where inputs to the hidden layer are now given by

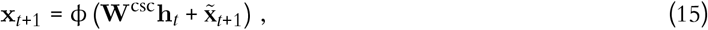

where 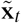 are the unmodified task inputs to the network that usually serve as **x**_*t*_. This setup allows for the photostimulation to propagate through the input weights since they now participate in a multi-synaptic recurrent loop, making them more similar to local recurrent weights. We initialize 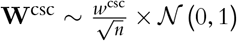. The size of *w*^*csc*^ strongly affects network dynamics through how much of an effect the feedback connections have on the input neuron activity. We quantify the size of this effect by comparing the relative size of the feedback connection inputs to the unmodified task inputs,

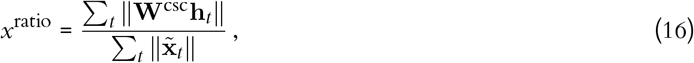

with *x*^ratio^ = 0 when *w*^csc^ = 0.

#### Preparatory model

To test whether learning-related changes in preparatory connectivity are sufficient to reshape BCI-relevant activity, we constructed a simplified firing-rate network model capturing the transition from preparatory to trial-period dynamics. The model was designed to reproduce both the baseline causal connectivity and learning-related changes in causal connectivity observed experimentally (Figs. 5, 6). Hidden activity evolved according to

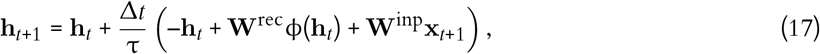

where **h**_*t*_ denotes population activity, Δ*t* = 0.01 s, τ = 0.1 s, and ϕ(*x*) = max(*x*, 0). Unlike the models described above, here the weight changes to the network are purely dictated by the observed changes (i.e. not optimized via gradient-descent).

The network comprised a pretrial population (10 neurons) generating low-dimensional preparatory activity, a downstream TRIAL population (10 neurons), five identical four-neuron modules that were initially untuned, and a single reward neuron. One neuron within one module was designated as the conditioned neuron (CN), and the BCI output was defined as its activity. Prior to learning, the CN lay outside the preparatory-to-trial pathway and exhibited weak trial responses (Fig. 7d).

Learning was implemented by modifying network connectivity to try to match the experimentally observed changes in causal connectivity. First, to reproduce the strengthened ‘pretrial →Δpretrial’ interactions, we increased connections from the pretrial population onto one neuron per initially untuned module, causing these neurons to acquire preparatory tuning and become Δpretrial neurons. This manipulation increased pretrial activity and enhanced the late trial-period response of the CN. However, this change alone did not reproduce the experimentally observed ‘Δpretrial →All’ connectivity pattern. To reproduce the experimentally observed ‘Δpretrial →All’ connectivity pattern (Fig. 7), we directly strengthened connections from Δpretrial neurons onto all downstream populations.

As an alternative implementation (Fig. S7), we instead strengthened connections from Δpretrial neurons back onto the pretrial population, thereby incorporating Δpretrial neurons into the preparatory subnetwork. Because pretrial neurons project broadly across the circuit, this modification caused Δpretrial neurons to effectively project broadly as well, reproducing the ‘Δpretrial →All’ connectivity pattern observed in the data. No synaptic weights were updated online; instead, learning was implemented by manually adjusting connectivity in a manner consistent with experimentally measured plasticity.

## Acknowledgments

This work was supported by the Paul G. Allen Foundation, Google DeepMind, NIH awards RF1DA055669 (SM, KA), R01EB029813 (SM) and R00MH121533 (MDG), and NSF award 2223725 (SM). We thank Ulises Pereira-Obilinovic, Lu Mi, Joseph Pemberton, and Fereshteh Lagzi for helpful comments and discussions. We also wish to thank the Allen Institute founder, Paul G. Allen, for his vision, encouragement, and support.

## Author contribution matrix

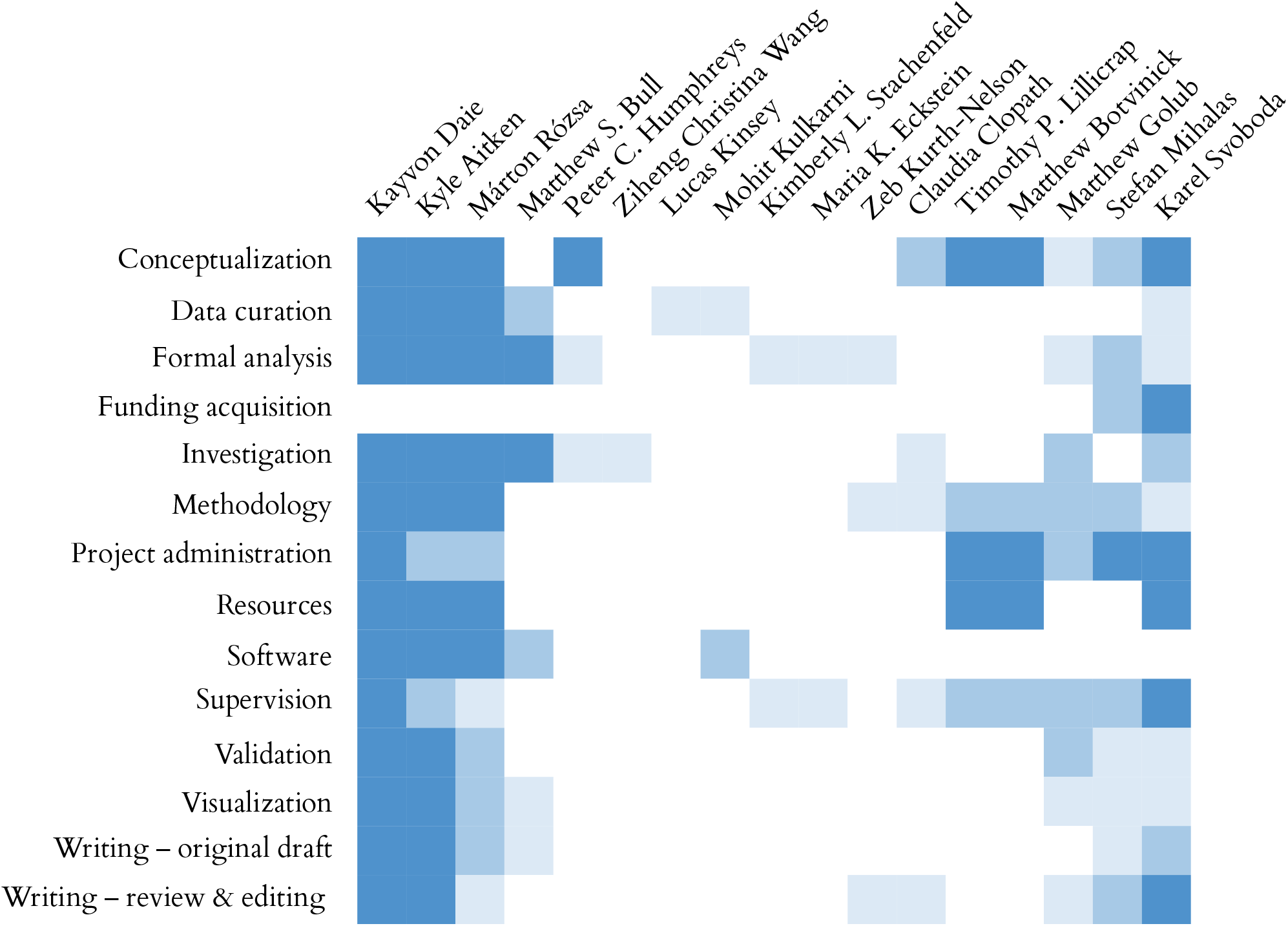

## Supplemental Figures

**Figure S1:**
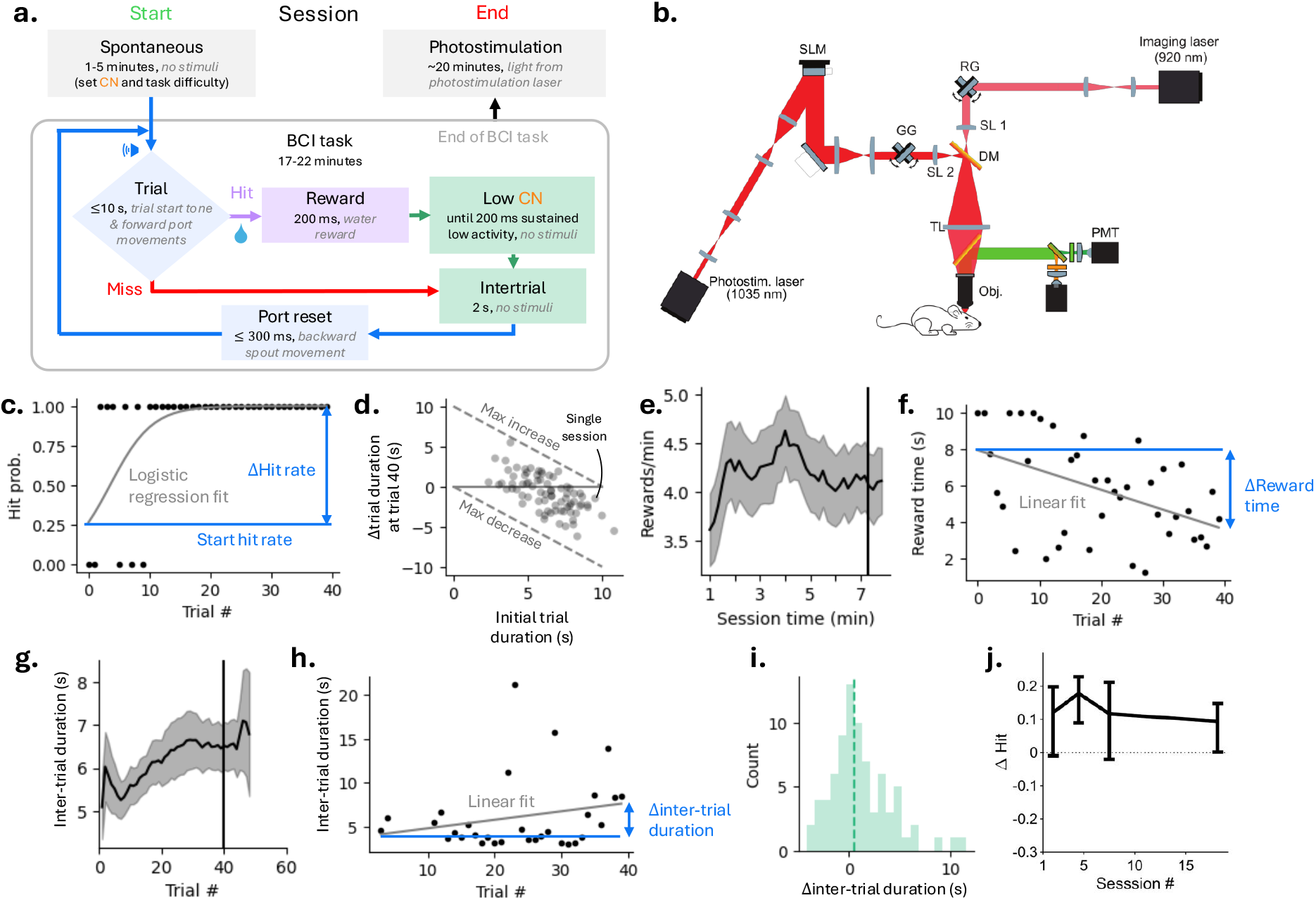
**(a)** Schematic of BCI trial structure. **(b)** Schematic of microscope. **(c)** Exemplar logistic regression fit on hit trial probability versus trial number, over first 40 trials. Same exemplar session shown in Fig. 1. Change in hit rate defined as difference in fit over first 40 trials. **(d)** Change in trial duration versus starting trial duration for all sessions. Dotted diagonal lines show maximum and minimum possible trial duration improvement given starting trial duration. **(e)** rewards per minute versus session time (mean over sessions, 95% bootstrap shading). Black line shows median time to 40 trials. **(f)** Linear regression fit to time to reward (from trial start) versus trial number for an exemplar session, over first 40 trials. Same exemplar session shown in Fig. 1. Change in reward time defined as difference in fit over first 40 trials. **(g)** Inter-trial duration, the time to next trial start after reward time, versus trial number (mean over sessions, 95% bootstrap shading). **(h)** Exemplar linear regression fit to inter-trial duration versus trial number for an exemplar session, over first 40 trials. Same exemplar session shown in Fig. 1. **(i)** Histogram of change in inter-trial durations over first 40 trials, dotted line shows median across sessions. **(j)** BCI performance (Δhit rate) versus number of sessions, showing negligible task meta-learning (mean, error bars bootstrapped 95% CI).

**Figure S2:**
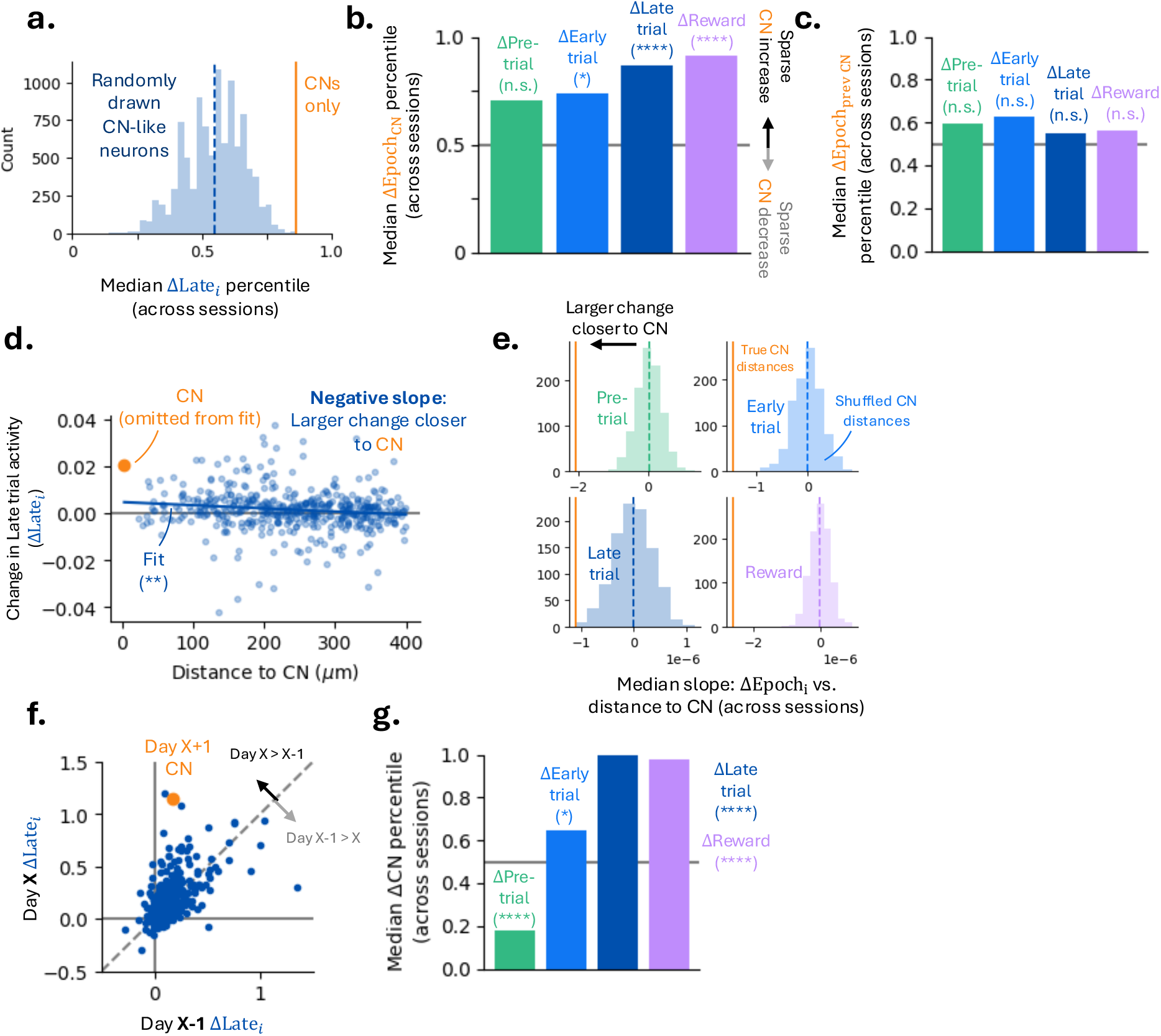
**(a)** Same as Fig. 2f, but random neurons are only drawn from neurons that would be considered candidate CN neurons (Methods). **(b)** Same as Fig. 2g, with random neurons drawn from CN-like candidates only. From left to right, *p* = 0.088, 0.088, < 10^−4^, < 10^−4^. **(c)** Same as Fig. 2g, with the CN from the previous day instead of the current day. **(d)** Change in late trial activity versus distance to the CN for an exemplar session. Linear fit, omitting CN, shown. Distance to CN truncated to 400 µm to minimize edge of FoV contributions. **(e)** Distribution of median slope of ΔEPOCH versus distance to the CN across sessions (e.g. as shown in (d)), for all 4 epochs. Distribution comes from shuffling distances to CN across neurons (10^3^ samples). Orange line shows median across unshuffled distance to CN. Unshuffled slopes consistently show negative trends, indicating larger changes closer to the CN. **(f)** Across session change of late trial, showing values for each neuron on Day X and Day X+1. **(g)** Same as Fig. 2g, where across session change, shown in (f), is used instead of within session change.

**Figure S3:**
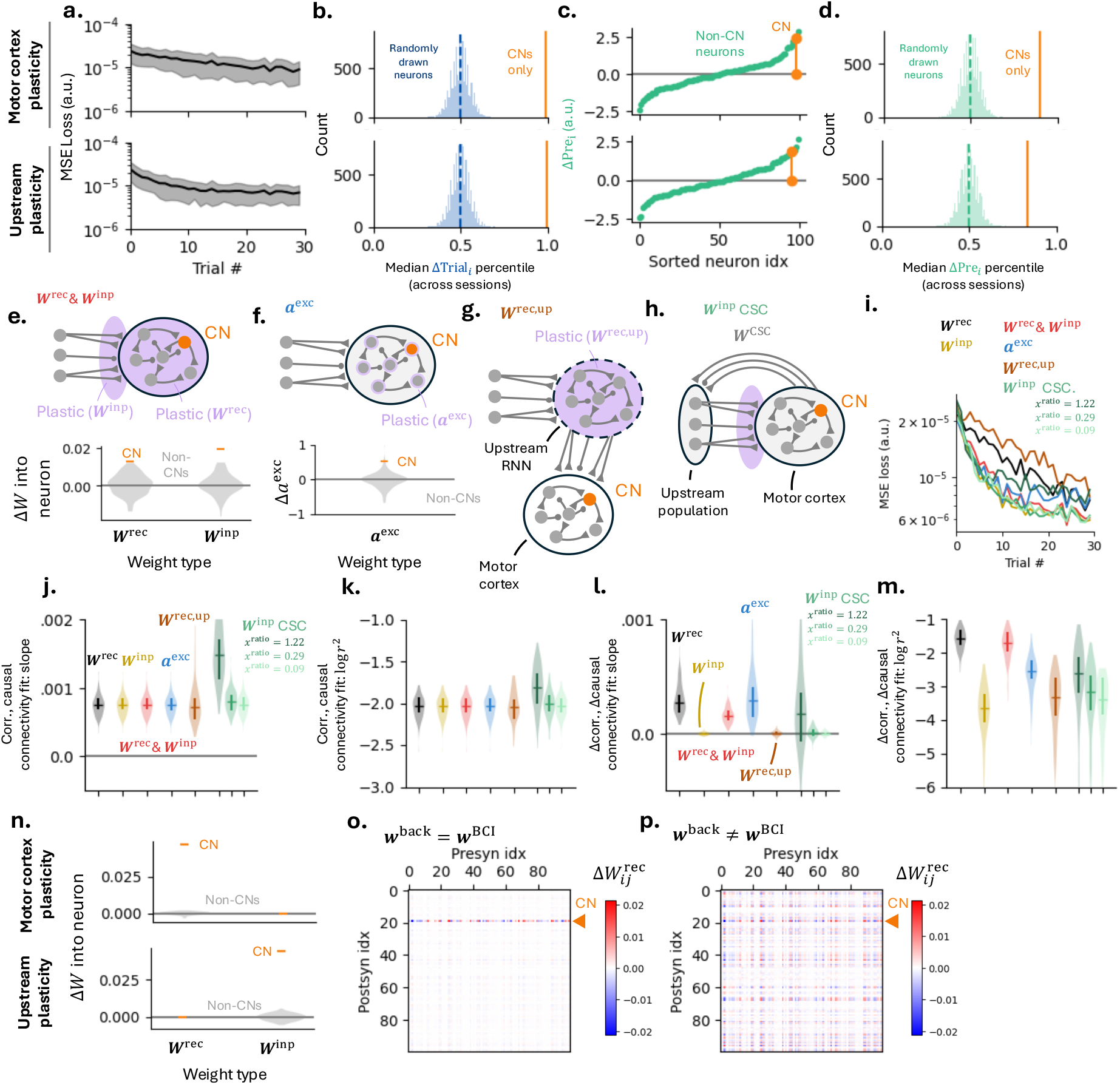
**[a-d]** Additional results from local MC (top) and upstream (bottom) plasticity models. Aggregate results averaged over 100 separate initializations. **(a)** MSE loss versus trial # (mean with standard deviation shading). **(b)** Distribution of median ΔTRIAL percentile across all sessions for randomly sampled neurons from each session, 10^4^ samples. Orange line shows median across the CN from all sessions. Model version of Fig. 2f. **(c)** Same as Fig. 3c, for pretrial. **(d)** Same as (b), for pretrial. **[e-m]** Alternative model comparisons. **(e)** Top: schematic of RNN model that learns via both MC and upstream plasticity (‘**W**^rec &^ **W**^inp^’). Bottom: exemplar weight changes from training. **(f)** Top: schematic of excitability trained RNN (‘**a**^exc^’). Bottom: exemplar excitability changes from training. **(g)** Schematic where upstream input is another RNN and its recurrent layer is trained (‘**W**^rec, up^’). **(h)** Schematic of RNN with additional feedback from hidden layer to input layer, potentially representing cortical-subcortical loops (‘**W**^inp^ CSC’). **(i)** Median loss functions across 100 initializations of each type of network. ‘**W**^inp^’ and ‘**W**^rec^’ are the upstream and MC plasticity models discussed in the main text. Distinct cortical-subcortical loop models shown for different connection strengths, quantified by *x*^ratio^ > 0, with larger values corresponding to stronger cortical-subcortical connections (Methods). **(j)** Correlation-causal connectivity fits performed analogously as Fig. 3f for each initialization. Distribution of slopes from fits across 100 initializations of each network type (dark line: median with 1st-3rd quartile bar). **(k)** Same as (j), but *r*^2^ of fits. **(l)** Same as (j), but Δcorrelation-Δcausal connectivity fits, analogous to Fig. 3g. **(m)** Same as (l), but *r*^2^ of fits. **[n-p]** Feedback misalignment of models (Methods). **(n)** Same as Fig. 3d, but for networks with feedback weights equal to BCI mask, **w**^back^ = **w**^BCI^. **(o)** Exemplar Δ**W**^rec^ over training for a network where **w**^back^ = **w**^BCI^. Weights with postsynaptic CN indicated. **(p)** Same as (o), but where **w**^back^ ≠ **w**^BCI^, representative of models used in main text.

**Figure S4:**
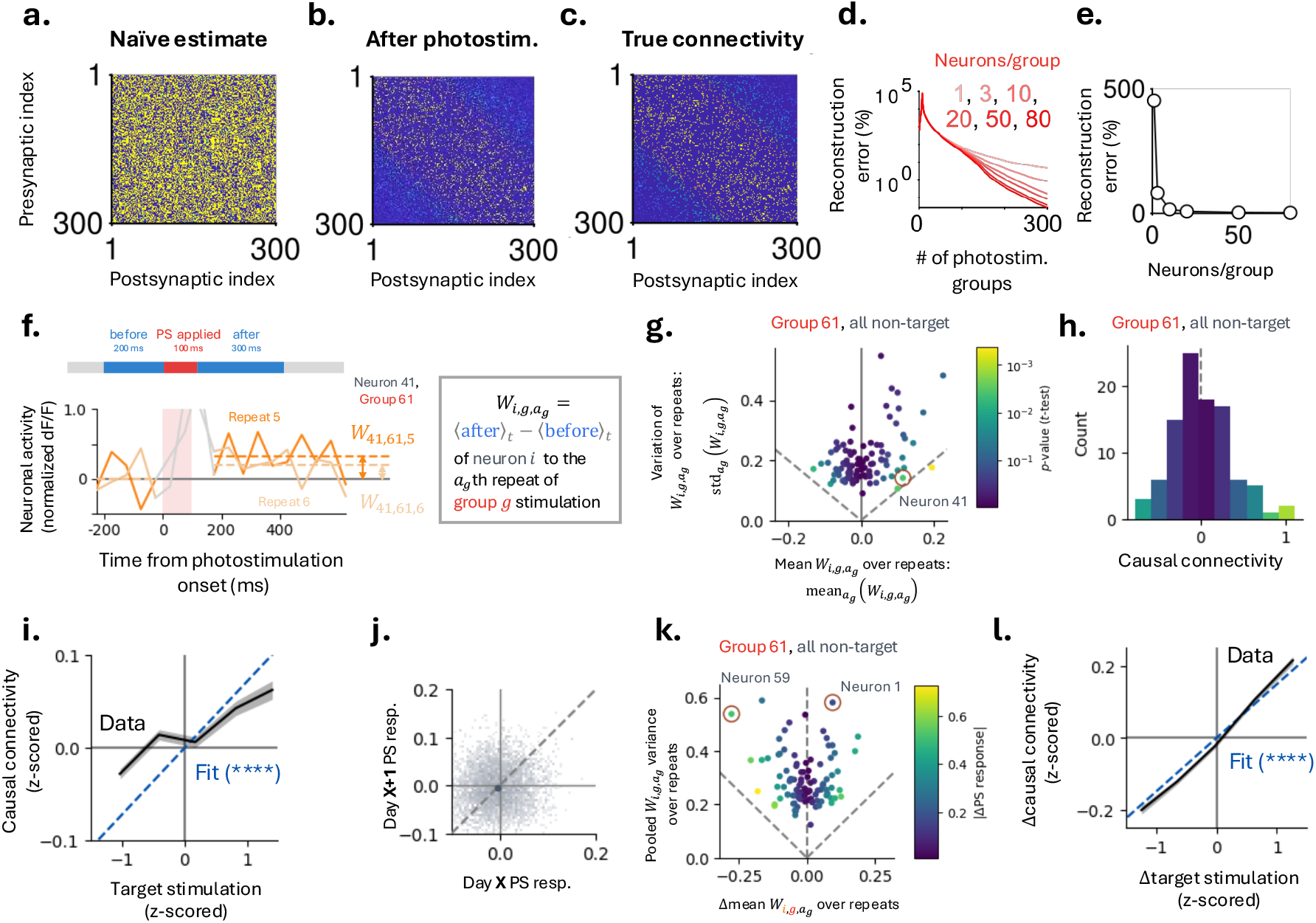
**[a-e]** Photostimulation reconstruction versus number of stimulated sites. **(a)** Estimate of reconstructed connectivity from spontaneous activity. **(b)** Same as (a), from spontaneous activity and 300 photostimuli with 80 neurons per photostimulation group. **(c)** True connectivity of estimates in (a) and (b). **(d)** Reconstruction error versus number of distinct photostimulation groups for 1, 3, 10, 20, and 80 neurons per stimulation group. **(e)** Reconstruction error versus number of neurons per group, for 300 photostimulation groups. **[f-i]** Causal connectivity. **(f)** Change in activity exemplar non-target neuron for two photostimulation repeats of same group (left). Definition of repeat response, which quantifies the photostimulation response to a given stimulation (right). **(g)** Mean repeat response versus standard deviation of repeat response, over repeats, for all non-target neurons of a given photostimulation group. Neurons colored by significance of response (*t*-test). **(h)** Causal connectivity of neurons shown in (g), colored by mean significance, same scale as (g). **(i)** Total target photostimulation versus non-target causal connectivity. **[j-l]** Change in causal connectivity. **(j)** Same as top of Fig. 4f, for non-target neurons. **(k)** Change in mean repeat response versus pooled repeat response variances, see Eq. (4). Neurons colored by Δcausal connectivity. **(l)** Change in total target photostimulation versus non-target Δcausal connectivity.

**Figure S5:**
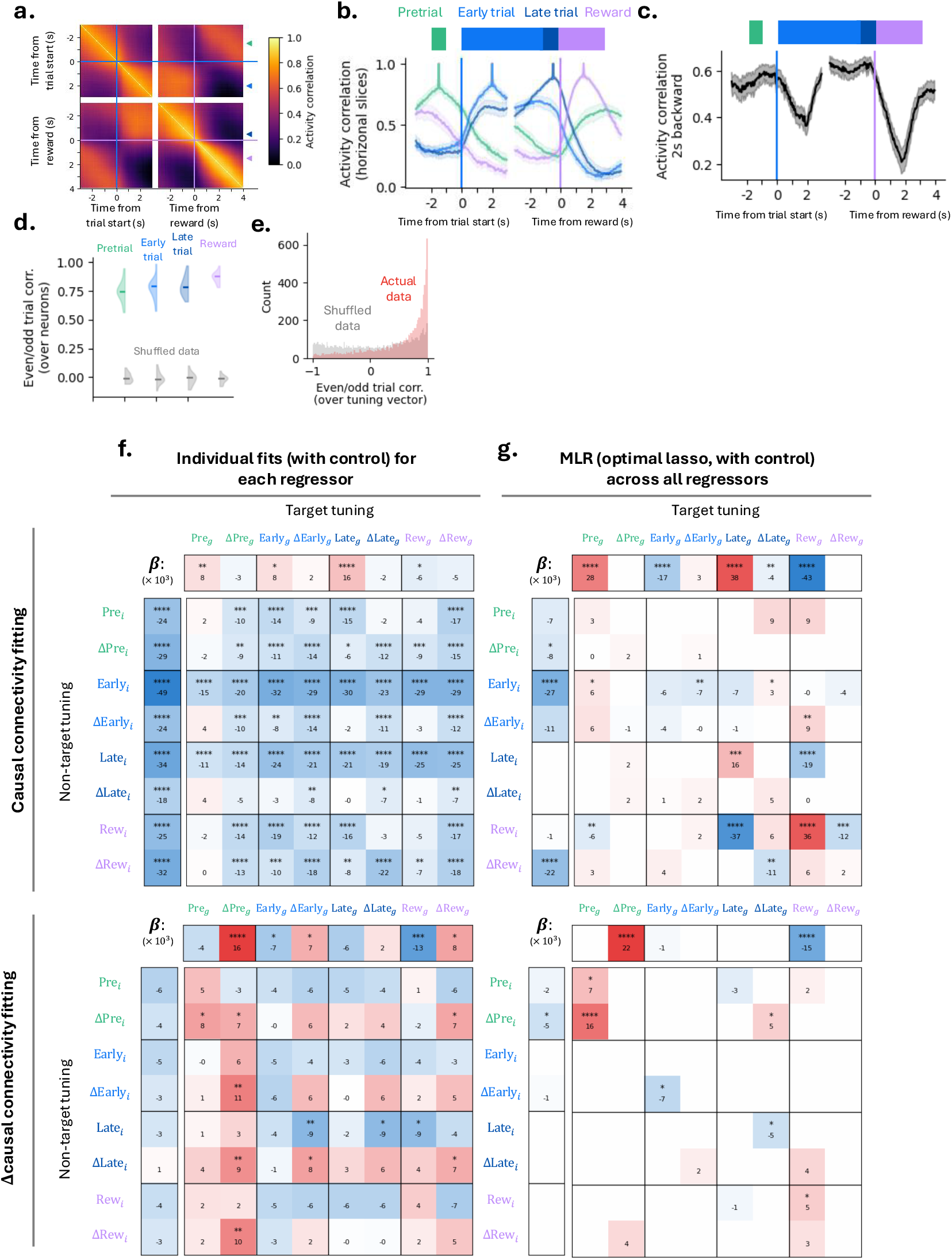
**(a)** Correlation of activity across task epochs, averaged across sessions. Horizontal slices shown in (b) denoted by triangles on right side. **(b)** Horizontal slices of similarity for exemplar times within each of the task epochs. **(c)** Activity correlation of current and 2 seconds backward showing variation in population activity versus time. **(d)** Correlation of neurons’ tuning when computed on only even/odd trials, computed over neurons for each task epoch. Grey shows data shuffled over neurons, solid line shows median over sessions. **(e)** Correlation of the tuning vector for each neuron when computed on only even/odd trials, computed over tack epochs for each neuron. **(f)** Fitting of causal connectivity (top) and Δcausal connectivity (bottom) using each individual regressor used in Figs. 5, 6. Unlike the MLR fits used in Figs. 5, 6, each fit only consists of the corresponding regressor and the control (Methods). **(g)** MLR fitting of causal connectivity (top) and Δcausal connectivity (bottom) using the eight-dimensional task tuning vectors of Fig. 6 (i.e. top is generalization of Fig. 5i, bottom is same as Fig. 6c for easy comparison).

**Figure S6:**
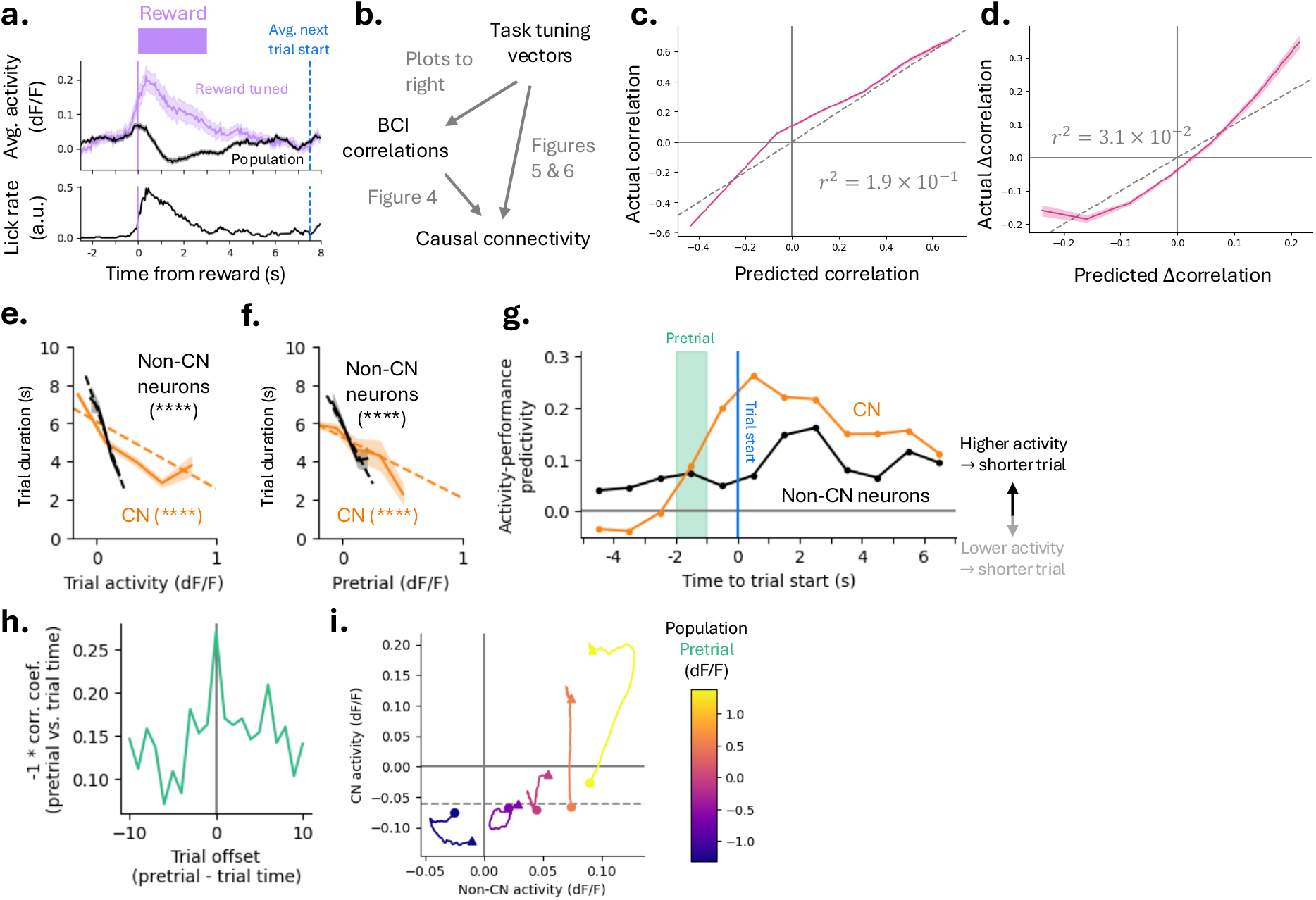
**(a)** Mean reward-aligned activity for reward responsive neurons and full population activity (top) and lick rate (bottom) as a function of time from reward. Shows population activity is modulated during reward consumption. **(b)** Relationship between the various routes of explaining causal connectivity and correlations of activity during BCI task used throughout the text. **(c)** MLR model fitting BCI correlations, uses same regressors as those determined from cross-validated lasso in Fig. 5i. **(d)** MLR model fitting ΔBCI correlations, uses same regressors as those determined from cross-validated lasso in Fig. 6c. **(e)** Trial duration as a function the CN (orange) and mean non-CN (black) trial activity, showing higher trial activity leads to shorter trial durations. Trial activity defined as the mean activity over entire trial (until timeout or reward). Solid lines show mean binned data, with s.e.m. shading. Dotted lines are linear regression fits. **(f)** Same as (e), but now as a function of pretrial activity. **(g)** Activity-performance predictivity, defined as the –sgn(slope) ∗*r*^2^ of the linear regression fits shown in (e) and (f), for 1 second bins aligned to trial start. Shows predictivity increases within trial, as expected, but remains high directly prior to trial start as well. Pretrial shaded region corresponds to dotted lines in (f). **(h)** How correlation of pretrial activity and trial duration changes as pretrial activity and trial duration are offset from one another. Shows correlation magnitude is highest for pretrial directly prior to said trial (i.e. 0 offset). **(i)** Evolution of non-CN activity and CN activity from low-CN time point to trial start, same binning shown in Fig. 6e.

**Figure S7:**
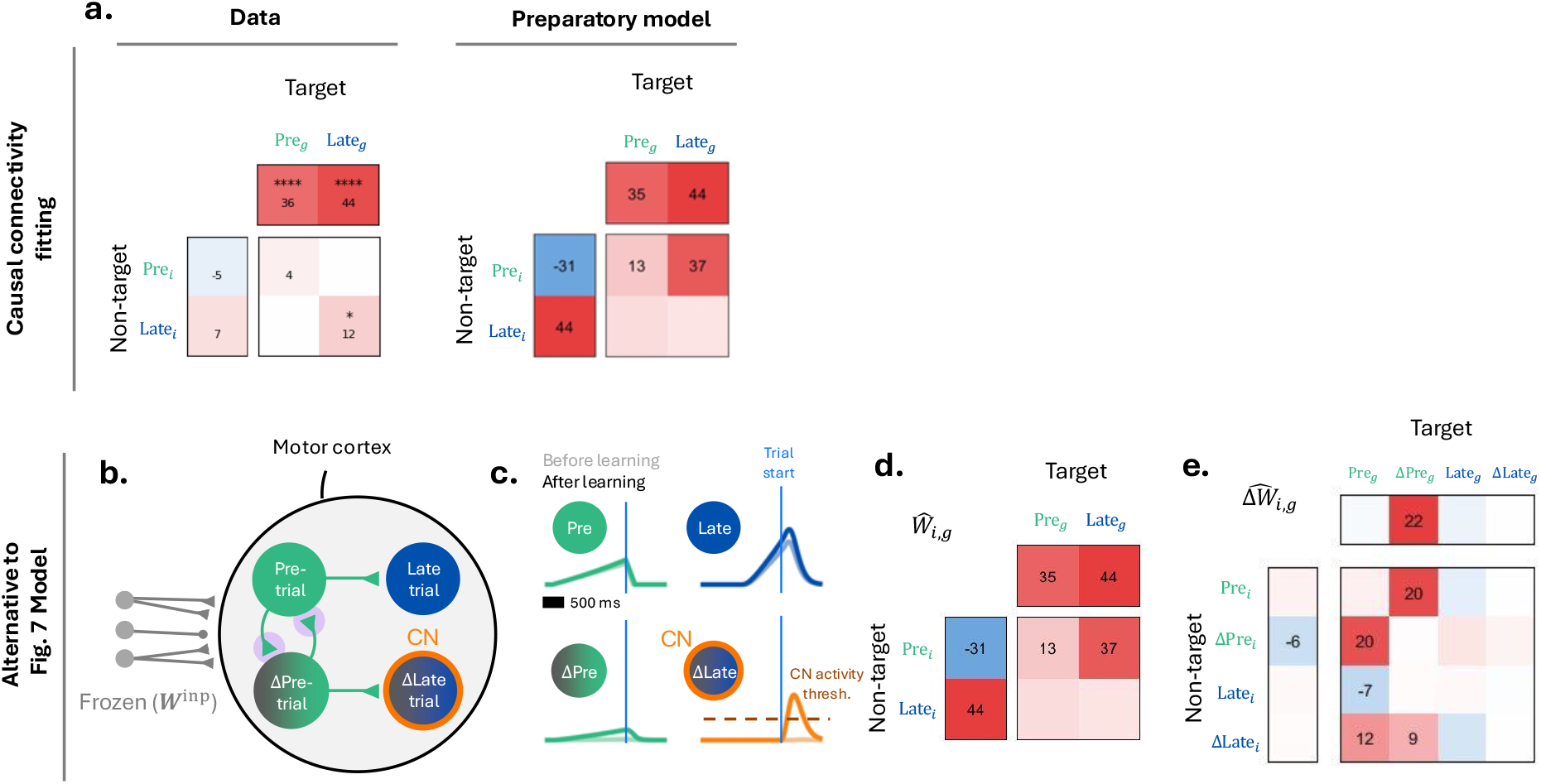
**(a)** Left: Subset of MLR coefficients fitting causal connectivity, shown in Fig. 5h, to be matched by the model. Right: MLR coefficients fitting model causal connectivity, indicating which task-tuning features predict connectivity (colored by size of coefficient, a.u.). **[b-e]** Same as Fig. 7, for an alternative weight change configuration. Schematic diagram of the preparatory network. Magenta circles indicate new connections that are added during learning. Activity of pretrial, Δpretrial, late trial, and Δlate trial neurons before (light) and after (dark) learning. **(d)** Same as (a, right), for alternative weight change configuration. **(e)** Δcausal connectivity matrix calculated using MLR as in Fig. 6c.

## Footnotes

1 In practice, it is possible to derive the perhaps more intuitive influence of a *single* target neuron on a non-target neuron. For example, one can take the Moore-Penrose inverse of photostimulation group membership matrix. This will only work for all neurons if the dimensionality of the target mask is equal to the number of neurons. In our case the upper bound of this dimensionality is the number of groups (100) which is always less than the number of neurons in the FoV (median: 481).

